# Murine GMP- and MDP-derived classical monocytes have distinct functions and fates

**DOI:** 10.1101/2023.07.29.551083

**Authors:** Sébastien Trzebanski, Jung-Seok Kim, Niss Larossi, Ayala Raanan, Daliya Kancheva, Jonathan Bastos, Montaser Haddad, Aryeh Solomon, Ehud Sivan, Dan Aizik, Jarmila Kralova, Mor Gross-Vered, Sigalit Boura-Halfon, Tsvee Lapidot, Ronen Alon, Kiavash Movahedi, Steffen Jung

## Abstract

Monocytes are short-lived myeloid immune cells that arise from adult hematopoiesis and circulate for a short time in the blood. They comprise two main subsets, in mice defined as classical Ly6C^high^ and non-classical Ly6C^low^ monocytes (CM, NCM). Recent fate mapping and transcriptomic analyses revealed that CM themselves are heterogeneous. Here, we report surface markers that allow segregation of murine GMP- and MDP-derived CM in the BM and blood. Functional characterization, including fate definition following adoptive cell transfer, established that GMP-Mo and MDP-Mo could equal rise to homeostatic CM progeny, such as NCM in blood and gut macrophages, but differentially seeded selected other tissues. Specifically, GMP-Mo and MDP-Mo gave rise to distinct interstitial lung macrophages, thus linking CM dichotomy to previously reported pulmonary macrophage heterogeneity. Collectively, we provide comprehensive evidence for the existence of two functionally distinct CM subsets in the mouse, which differentially contribute to peripheral tissue macrophage populations in homeostasis and following challenge. Our findings are indicative of impact of monocyte ontogeny on *in situ* differentiation.

## Introduction

Monocytes are circulating short-lived myeloid immune cells that can promote inflammation and serve as precursors of tissue-resident macrophages (MΦ) ^1^. Monocytes arise from committed progenitors in the bone marrow (BM) ^2,3^ and are released into the blood stream upon maturation. Under homeostatic conditions, monocytes replenish selected tissue MΦ compartments to various extent. This includes barrier organs, such as the gut and skin, but also tissues that experience mechanical stress, like the heart ^4^.

In humans, mice, and other mammals, two main subsets of blood monocytes have been identified ^5–8^. Murine monocytes have been subdivided into Ly6C^high^ CCR2^+^ ‘inflammatory’ or classical monocytes (CM) and Ly6C^low^ CCR2^−^ ‘patrolling’ or non-classical monocytes (NCM) ^6,7,9^. Apart from distinct surface marker profiles and transcriptomes ^10^, monocyte subsets differ, as suggested by their names, also in function. CM are generated in the BM and have short circulation half-lives of about a day in both mice and humans ^11,12^. CM are major contributors to tissue resident MΦ populations, both in homeostasis and inflammation ^1,11,13^. Within tissues, CM give rise to monocyte-derived MΦ and, potentially, CD209a^+^ monocyte-derived dendritic cells (MoDC) ^14–17^. NCM, on the other hand, arise in the circulation from CM in a Notch-dependent manner ^18–20^. NCM patrol the wall of blood vessels ^9,21^, depend on Cx_3_cr1 for survival ^22^, are longer-lived than CM ^11,12^, and can be considered vasculature-resident MΦ. Further, NCM have been reported to give rise to tissue-resident cells ^23–25^, although their precursor function is less well established.

Recent studies have revealed further complexity in BM monopoiesis ^26,27^. Specifically, adoptive cell transfer and lineage tracing studies suggest a developmental CM bifurcation. CM that arise from granulocyte and macrophage progenitors (GMP) display transcripts encoding neutrophil granule proteins, such as elastase (*Elane)*, myeloperoxidase (*Mpo*), and chitinase-like protein 3 (*Chil3)* ^26,27^. Conversely, CM emerging from monocyte and DC precursors (MDP) ^28^ harbor signatures enriched for transcripts related to MHC-II expression and encoding classical DC markers, such as *Cd209a* ^26,27^. Interestingly, and suggesting discrete functional roles of these CM subsets, challenges result in selective expansion of these GMP-Mo and MDP-Mo populations ^26^. However, CM heterogeneity remains largely defined by transcriptomics and functional contributions and fates of GMP-Mo and MDP-Mo upon their recruitment into tissues remain thus to be elucidated.

Here, we report surface markers that allow the discrimination of murine GMP-Mo and MDP-Mo as CD177^+^ and CD319^+^ CM, respectively, enabling us to perform a comprehensive analysis of *in vitro* and *in vivo* features of these cells. Corroborating earlier work, mice challenged with different microbial stimuli exhibited distinct GMP-Mo and MDP-Mo dynamics. Further, classical *in vitro* assays for neutrophil-related activities established unique functional potential of GMP-Mo. Competitive adoptive transfers of the two CM populations into MΦ-depleted animals revealed overlapping and distinct contributions of GMP-Mo and MDP-Mo to peripheral tissue-resident MΦ, including the gut, lung, and meninges. Likewise, also in thorax-shielded animals GMP and MDP gave rise to distinct pulmonary MΦ populations. Collectively, we link CM dichotomy to heterogeneity of pulmonary MΦ and establish differential contributions of GMP-Mo and MDP-Mo to the lung in health and following viral challenge.

## Results

### Identification of surface markers discriminating murine classical monocyte subsets

Emerging evidence suggests discrete subpopulations among CM that arise from GMP and MDP, respectively ^8,26,27^. GMP-Mo have been reported to display a neutrophil-like gene signature, while MDP-Mo were noted to bear DC expression features. However, except for a recent study ^29^, CM heterogeneity has not been established beyond mere transcriptomics.

In search for surface markers that will allow to investigate functional aspects of the CM subsets in various physiological settings, we performed a comprehensive CITE-seq screen on whole BM isolated from C57BL/6 wild type (wt) animals (**Fig 1A**). All myeloid immune subsets and their precursors were well represented in our data set (**Fig 1B, Suppl Fig 1A,B,C**). Anti-Ly6C and -CD115 TotalSeq antibodies reliably identified *Csf1r*^+^ *Fcgr3*^+^ monocytes and their precursors (**Fig 1C**). Focusing on mature monocytes and nearby clustering classical DC (cDC), we identified a small cluster of Ly6C^+^ CD115^+^ *Fcgr3*^+^ monocytes devoid of the CM marker CD62L (encoded by *Sell*) ^6,7^ (**Fig 1D**). Comparison of CD62L^+^ and CD62L^−^ CM clusters highlighted enriched expression of neutrophil-related genes, such as *Chil3*, *Mmp8,* and *Slpi* among CD62L^+^ CM. Conversely, MHC-II transcripts were upregulated in the CD62L^−^ subset (**Fig 1D,E**). Interestingly, and despite their high expression of MHC-II transcripts, these cells were otherwise transcriptionally and phenotypically distinct from cDC, suggesting that they might be previously reported DC-like monocytes (**Suppl Fig 1D**).

**Figure 1:**
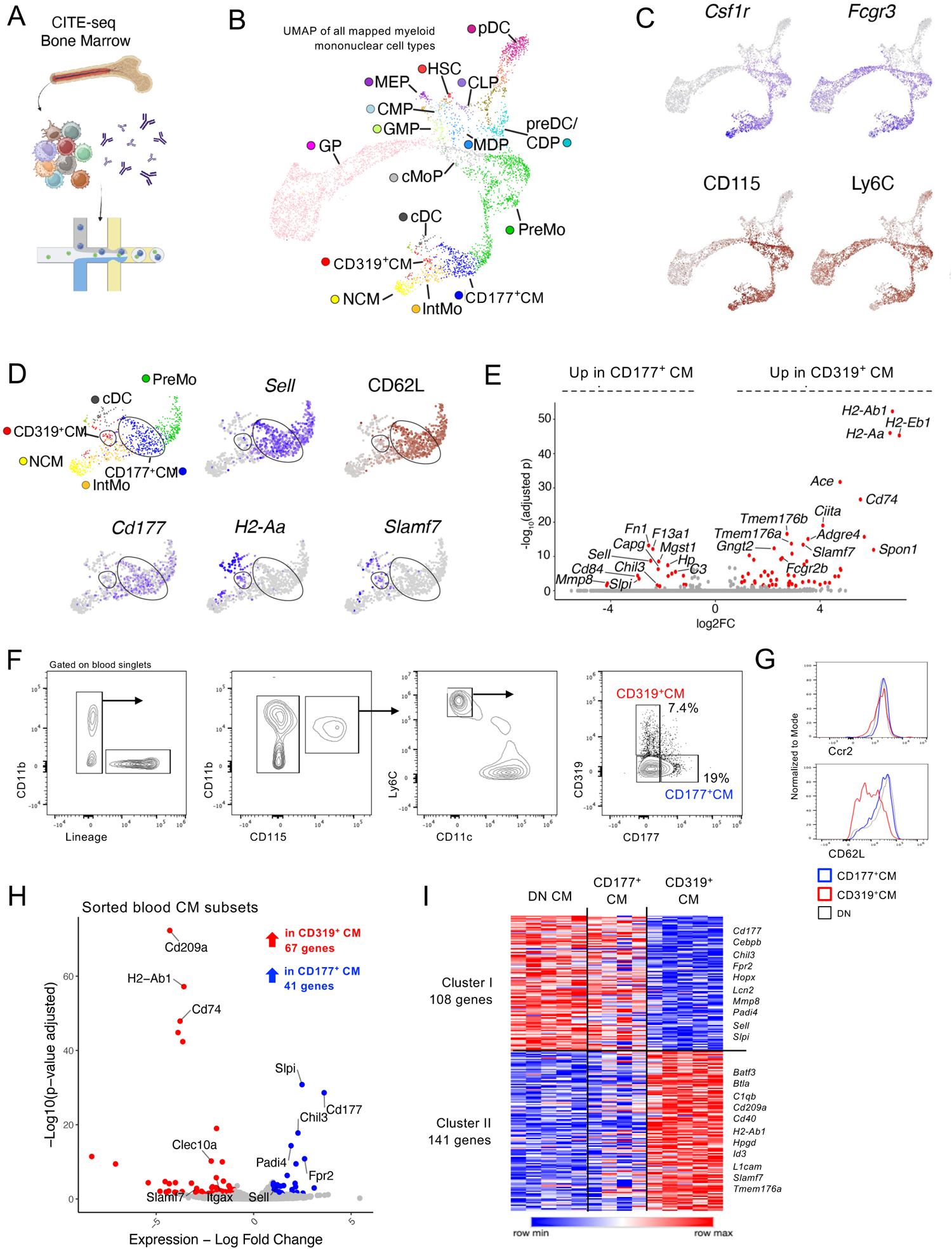
Identification and characterization of CM subsets by scRNAseq. (A) Experimental scheme for the CITE-seq experiment. (B) Curated UMAP plot of the BM monocytes, DCs and their precursors identified by CITE-seq. (C) UMAP plots showing the protein expression of Ly6C and CD115 as identified by CITE-seq antibodies, and Csf1r and *Fcgr3* gene expression in the dataset from B. (D) UMAP plot, visualizing the mature monocyte and cDC subsets, and the gene or protein expression of selected markers. (E) Differentially expressed genes between the CD177^+^ CM and CD319^+^ CM clusters identified by CITE-seq. (F) Gating on blood monocytes identifying CD177 and CD319 as markers for monocyte subsets. (n=12 from 8 independent experiments) (G) Surface expression of canonical monocyte markers on CD177^+^ Ly6C^high^ CM (blue), CD319^+^ Ly6C^high^ CM (red), and CD177^−^CD319^−^ Ly6C^high^ CM (black). (n=6 from 2-3 independent experiments) (H) Volcano plot showing differentially expressed genes between CD177^+^ CM (blue dots) and CD319^+^ CM (red dots) from sorted blood cells subjected to bulk RNAseq. CM were defined as Lin^−^ CD11b^+^ CD115^+^ Cx_3_cr1^GFP+^ Ly6C^high^ CD11c^−^ cells. (n=4-5) (I) Heatmap showing differentially expressed genes between DN (CD177^−^CD319^−^ CM), CD177^+^ CM, and CD319^+^ CM from sorted blood cells. (n=4-5)

Closer examination of the list of differentially expressed genes (DEG) between the CD62L^+^ and CD62L^−^ monocyte clusters revealed *Slamf7* (encoding CD319) restricted to CD62L^−^ cells (**Fig 1D,E**). Conversely, *Cd177* expression was rather unique to CD62L^+^ cells. Indeed, combined staining for CD319 (clone 4G2) and CD177 (clone Y127) provided segregation of two Ly6C^high^ CM subsets by flow cytometry (**Fig 1F**). Moreover, also an independent LEGENDscreen approach on total blood cells revealed CD319 to yield sufficient segregation of CM when plotted against CD177 (**Suppl Fig 1F**).

Phenotypic characterization of CD177^+^ and CD319^+^ blood CM by flow cytometry confirmed equal levels of the Ccr2 chemokine receptor on the surface of both cells, as well as lower CD62L expression on CD319^+^ CM (**Fig 1G**). CD177^+^ CM displayed a shift for CD88a and Ly6C, while CD319^+^ CM showed higher expression of MHC-II, the chemokine receptor Cx_3_cr1, and the Fc receptor CD64 (**Suppl Fig 1G**). Importantly and underlining the robustness of our finding, differential CD177 and CD319 expression on blood CM could be independently confirmed in another animal facility (**Suppl Fig 1H**). Lastly, CD177^+^ but not CD319^+^ blood CM expressed higher levels of CD157 (Bst1), in line with a recent report identifying CD157 and CD88a as markers of neutrophil-like GMP-Mo ^29^ (**Suppl Fig 1I**).

Also DC precursors were reported to display a number of classical CM surface markers, including CD115 and Ly6C ^30^. Therefore, we probed for a potential contamination of the CD319^+^CD11c^−^ CM gate with CD11c^+^MHC-II^−^ pre-cDC by analysis of Flt3 expression and intracellular staining for the cDC lineage-defining transcription factor (TF) Zbtb46 ^31,32^ (**Suppl Fig 1J**). Neither Zbtb46 nor Flt3 were found to be expressed by CD319^+^ CM (**Suppl Fig 1K**).

Finally, we sorted CD319^+^, CD177^+^, and CD319^−^CD177^−^ blood CM to purity and subjected them to bulk RNAseq (**Suppl Fig 1L**). Indeed, and akin to BM CM subsets (**Fig 1E**), CD177^+^ and CD319^+^ CM differed in the expression of neutrophil and DC gene modules, respectively (**Fig 1H,I**, **Suppl Fig 1M,N**). Further, elevated *Clec10a* and *Cd209a* transcripts in CD319^+^ CM correlated with higher surface protein expression (**Suppl Fig 1O**). Of note, CD319^−^ CD177^+^ and CD319^−^ CD177^−^ blood CM subsets were transcriptionally quasi-identical and, thus, most likely constitute the same cell type (**Fig 1I**).

In conclusion, we have identified CD177 and CD319 as surface markers delineating CM heterogeneity in line with previous reports. CD177^+^ CM were enriched in neutrophil-related gene modules, while CD319^+^ CM expressed high levels of *H2-Ab1* reminiscent of DC. Collectively, these data suggest that CD177^+^ and CD319^+^ CM are derivatives of GMP and MDP BM precursors.

### CD319^+^ CM predominantly arise from MDP

In their seminal study, Goodridge and colleagues proposed that the two CM subsets derive from GMP and MDP, respectively ^26^. Accordingly, also CD177^+^ CM would be expected to arise mostly from GMP, whereas CD319^+^ CM should be progeny of MDP. First, we adopted a fate-mapping strategy and crossed *Ms4a3^Cre^:R26^LSL-TdTom^* animals, which allow fate mapping of GMP ^13^, to *Cx3cr1^Gfp^* animals, labeling all monocytes. As previously reported ^13^, a small fraction of blood monocytes remains unlabelled in these double-reporter (DR) mice, likely constituting MDP-derived cells (**Fig 2A**). Interestingly, these unlabeled cells showed robust, albeit not exclusive, expression of CD319, as well as higher Cx_3_cr1 expression levels, akin to blood CD319^+^ CM (**Fig 2A, Suppl Fig 1G**).

**Figure 2:**
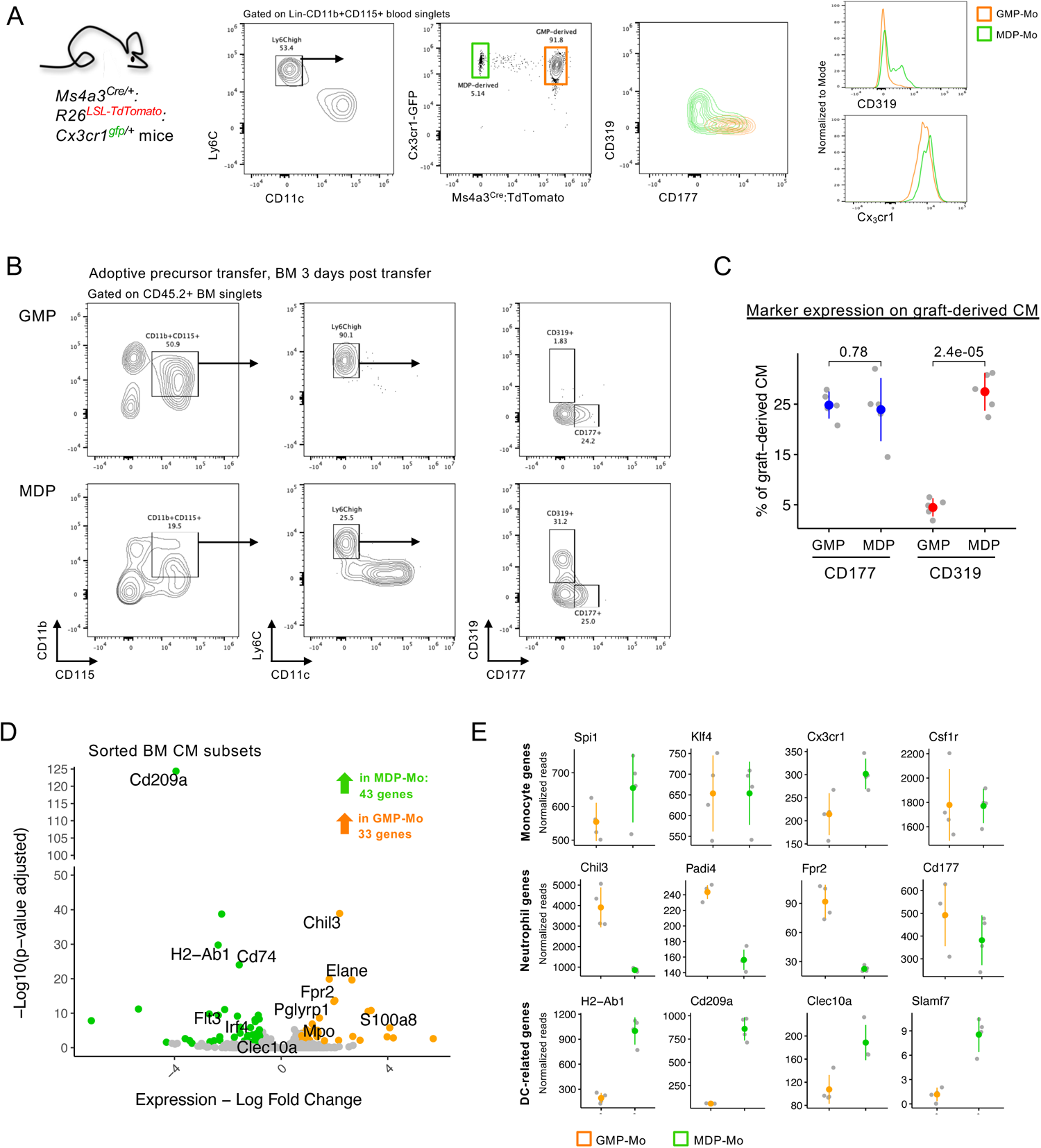
CD319^+^ CM predominantly arise from MDP. (A) Flow cytometry analysis of blood monocytes from *Ms4a3^Cre^:R26^LSL-TdTom^:Cx3cr1^Gfp^* mice. (n=6 from 2 independent experiments) (B) Flow cytometry analysis of adoptive precursor transfer experiments (CD45.2 > CD45.1). BM of recipients was analyzed on day 3 after transfer. (n=5-6 from 2 independent experiments) (C) Statistical analysis of CD177^+^ and CD319^+^ CM derived from adoptively transferred GMP or MDP. (n=5-6 from 2 independent experiments) (D) Volcano plot of differentially expressed genes among *Ms3a4^Cre^:R26^LSL-TdTom^-:Cx3cr1^Gfp^* (MDP-derived) monocytes and *Ms4a3^Cre^:R26^LSL-TdTom^+:Cx3cr1^Gfp^*(GMP-derived) monocytes sorted from BM according to their TdTomato and GFP expression, respectively. CM were defined as CD11b^+^ CD115^+^ CD11c^−^ MHC-II^−^ Ly6C^high^ cells (n=4) (E) Scatter plots of selected genes among BM GMP- and MDP-derived CM. *Chil3*, *Fpr2*, *H2-Ab1*, *Cd209a*, and *Clec10a* were differentially expressed. (n=4)

Next, we performed adoptive precursor transfers to directly link CD177 and CD319 expression to GMP-Mo and MDP-Mo, respectively (**Suppl Fig 2A**). Indeed, GMP gave rise to CD177^+^ CM but much less so to CD319^+^ CM (**Fig 2B**). On the other hand, one third of MDP-derived Ly6C^high^ CD11c^−^ CM expressed CD319 (**Fig 2B,C**). The abundance of CD177-expressing cells was similar among GMP- and MDP-derived CM (**Fig 2B,C**).

Finally, we isolated TdTom^+^ GFP^+^ and TdTom^−^ GFP^+^ CM from BM of *Ms4a3^Cre^:R26^LSL-TdTom^:Cx3cr1^Gfp^* double-reporter mice, further enriched for lack of CD11c and MHC-II expression to exclude cDC and their precursors (**Suppl Fig 2B**), and subjected them to bulk RNAseq analysis. In line with the proposed developmental scheme ^13,26^, TdTom^−^ GFP^+^ BM monocytes expressed higher levels of DC-related transcripts, such as *H2-Ab1* and *Cd209a* (**Fig 2D,E**). Likewise, TdTom^+^ GFP^+^ BM monocytes displayed a neutrophil-like gene expression signature, including *Chil3* and *Elane* (**Fig 2D,E, Suppl Fig 2C**). Of note, *Slamf7,* encoding CD319, was lowly expressed in BM monocytes (**Fig 2E**). This might be due to the generous CD11b sorting gate we used (**Suppl Fig 2B**), which most likely included immature CD11b^int^ transitional pre-monocytes ^33^. The latter could also explain the expression of *Flt3* and neutrophil granule proteins, such as *Mpo* and *Ngp*, which was absent from blood monocytes (**Fig 2D**).

In conclusion, we show through two independent approaches that CD319^+^ CM arise mainly, albeit not exclusively, from MDP; conversely, the vast majority of these CD177^+^ CM derives from GMP.

### Differential response of GMP-Mo and MDP-Mo to microbial stimuli

The CM compartment dynamically responds to challenges including exercise, metabolic alterations, and pathogen encounter ^34,35^. Under conditions of parasitic and bacterial inflammation associated with IFNγ exposure, CM have been shown to upregulate expression of the stem cell marker Sca1 (Ly6A) and concomitantly downmodulate the monocyte/ MΦ lineage marker Cx_3_cr1 ^36,37^.

GMP-Mo and MDP-Mo were reported to differentially expand in animals challenged with LPS and CpG ^26^. In line with these findings, exposure of wt C57BL/6 animals to LPS (2.5 μg/g, i.p. or i.v.) resulted by day 1 in an increase in CD177^+^ cells while MDP-Mo remained proportionally unaffected (**Fig 3A,B, Suppl Fig 3A**). Surprisingly, Cx_3_cr1 surface expression was selectively lost on GMP-Mo, whereas MDP-Mo retained closer-to-homeostatic Cx_3_cr1 levels (**Suppl Fig 3B**). Conversely, MDP-Mo and CD319^−^ CD177^−^ double-negative (DN) GMP-Mo, but not CD177^+^ GMP-Mo upregulated Sca1 as well as other previously reported MDP-Mo markers, such as CD64, CD11c, and CD209a (**Suppl Fig 3B**).

**Figure 3:**
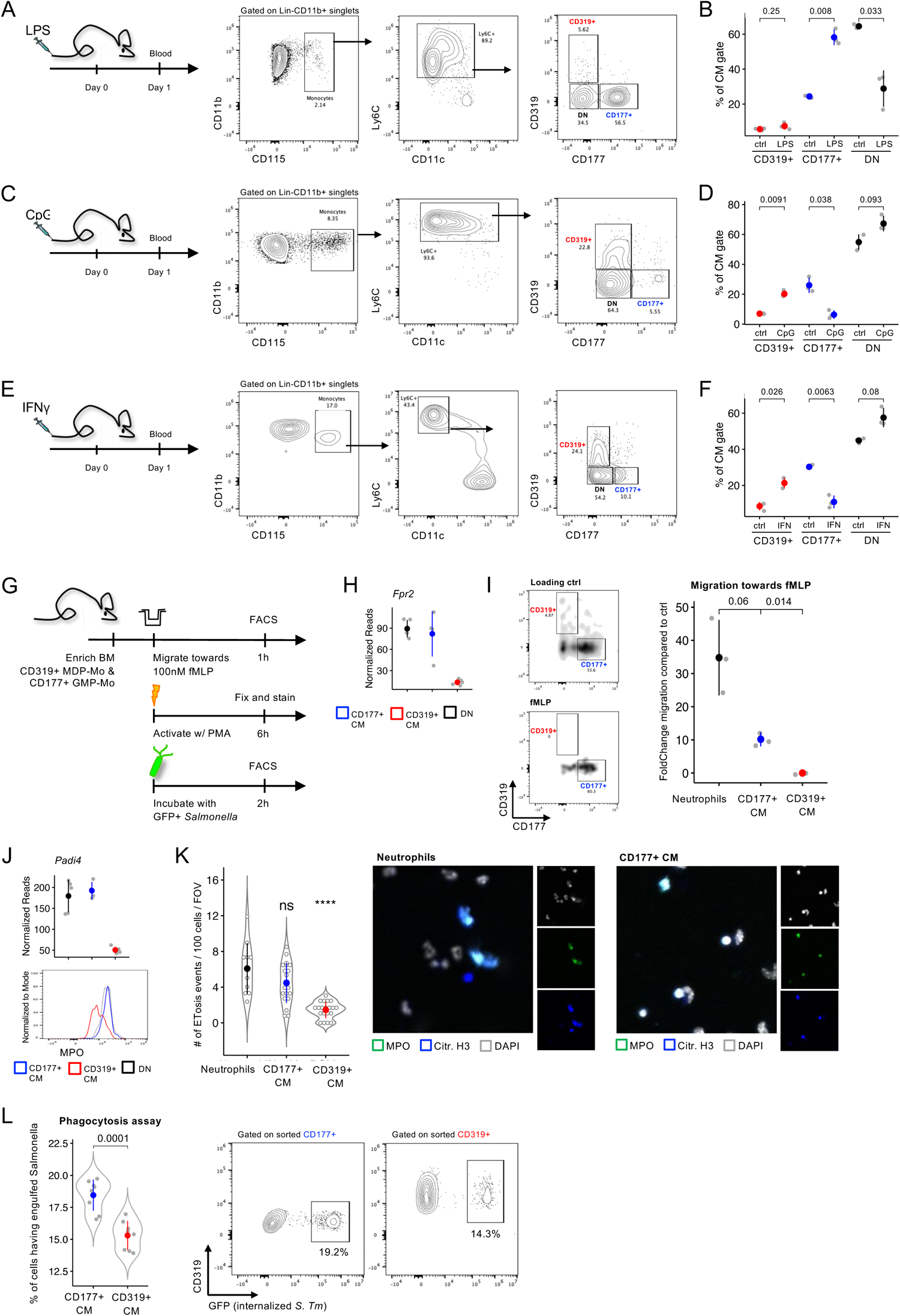
*In vivo* and *in vitro* characterization of GMP-Mo and MDP-Mo. (A) Experimental scheme and blood analysis of LPS-treated mice one day after treatment. (n=9 from 3 independent experiments) (B) Quantification of changes in the CM compartment after LPS treatment. (n=9 from 3 independent experiments) (C) Experimental scheme and blood analysis of CpG-treated mice one day after treatment. (n=6 from 2 independent experiments) (D) Quantification of changes in the CM compartment after CpG treatment. (n=6 from 2 independent experiments) (E) Experimental scheme and blood analysis of rIFNγ-treated mice one day after treatment. (n=6 from 2 independent experiments) (F) Quantification of changes in the CM compartment after rIFNγ treatment. (n=6 from 2 independent experiments) (G) Schematic of performed *in vitro* assays. (H) Transcriptional expression of the formyl-methionine peptide receptor 2 (Fpr2) among CM subsets. (n=4-5) (I) Quantification of migration of BM monocytes and neutrophils towards fMLP (100 nM) compared to control (DMSO), as assessed by flow cytometry. (n=6 from 2 independent experiments) (J) Transcriptional expression of the protein amine deiminase 4 (Padi4) protein among CM subsets and intracellular staining for myeloperoxidase (MPO) among CM subsets by flow cytometry. (n=4-5 for transcriptomics, n=6 from 2 independent experiments for MPO staining) (K) Quantification and microscopy images of the initiation of ETosis events as evidenced by co-staining for DNA, MPO, and citrullinated histone 3 (Citr. H3) as assessed by microscopy. (n=30-40 FOV of each cell type from 2 independent experiments) (L) Quantification and representative flow cytometry plots of phagocytosis of GFP-expressing *Salmonella Typhimurium* by GMP-Mo and MDP-Mo. (n=13 from 2 independent experiments)

The TLR9 agonist CpG, in combination with the cationic lipid DOTAP enhancing nucleic acid delivery, was reported to expand MDP-derived monocytes ^26^. Accordingly, administration of CpG + DOTAP (5 μg and 25 μg/mouse, respectively) increased the representation of CD319^+^ MDP-Mo within the blood Ly6C^+^ CM gate at the expense of CD177^+^ GMP-Mo (**Fig 3C,D, Suppl Fig 3C**). Notably, Sca1 was strongly upregulated on all CM subsets while Cx_3_cr1 expression remained unaltered (**Suppl Fig 3D**). In contrast to LPS-treated mice, and testifying monocyte plasticity, CD177^+^ GMP-Mo upregulated the CD319^+^ MDP-Mo markers CD64, CD11c, and CD209a following CpG exposure (**Suppl Fig 3D**).

Infection-associated IFNγ prompts the emergence of Sca1^+^ monocytes ^36,37^. Indeed, akin to CpG challenge, CD319^+^ MDP-Mo were significantly expanded at the expense of CD177^+^ GMP-Mo in the CM gate of mice given a one-time injection of 5 μg recombinant IFNγ (rIFNγ) (**Fig 3E,F, Suppl Fig 3E**). Cx_3_cr1 expression was not altered among subsets, while Sca1 was higher expressed on CD319^+^ MDP-Mo (**Suppl Fig 3F**). CD319^−^ CM modestly upregulated MHC-II, whereas CD319^+^ MDP-Mo expressed high levels of both MHC-II and CD11c (**Suppl Fig 3F**). Transcriptome analysis of sorted Sca1^+^ and Sca1^−^ CM retrieved from blood of animals that underwent repetitive rIFNγ challenges (**Suppl Fig 3G**) revealed that Sca1^+^ CM expressed *Slamf7* (encoding CD319), whereas Sca1^−^ CM displayed higher expression of *Cd177* and *Fpr2* (**Suppl Fig 3H,I,J**). This suggests that distinct CD177^+^ GMP-Mo and CD319^+^ MDP-Mo signatures are retained, at least on the transcriptional level, following challenges.

Collectively, and as previously described for GMP- and MDP-derived BM monocytes ^26^, GMP-Mo and MDP-Mo differentially expand following exposure to microbial stimuli, likely as a result of altered monopoiesis.

### GMP-Mo but not MDP-Mo display neutrophil-like functions *in vitro*

Neutrophils are first-responders at sites of sterile and non-sterile injury, where they function as phagocytes and neutralize extracellular pathogens ^38–40^. To investigate if neutrophil-like GMP-Mo share, beyond their transcriptomic overlap, functional neutrophil hallmarks, we performed a series of *in vitro* assays (**Fig 3G**).

N-formylpeptides are cleavage products of bacterial and mitochondrial proteins that act as potent neutrophil chemoattractants ^41^. To test for activity of N-formylpeptide receptor 2 (Fpr2) which was robustly and differentially expressed in CD177^+^ GMP-Mo (**Fig 3H**), we magnetically enriched CD115^+^ BM monocytes and subjected them to a migration assay towards 100 nM fMLP or carrier control (**Fig 3I, Suppl Fig 3K,L**). Analysis of migration towards fMLP-supplied medium revealed that CD177^+^ GMP-Mo displayed a 10-fold enrichment compared to the control condition, while CD319^+^ MDP-Mo did not migrate at all in this assay (**Fig 3I**).

A hallmark of neutrophil activation is the formation of extracellular traps (ET) that are linked to granule protein expression, including MPO, as well as histone citrunillation and deamination ^38^. CD177^+^ GMP-Mo, but not CD319^+^ MDP-Mo, expressed one of the key factors required for chromatin de-condensation, the protein-arginine deiminase type-4 (Padi4) as well as high levels of intracellular MPO ^42^ (**Fig 3J**). To test whether Padi4 endows GMP-Mo with potential to form ET, CD177^+^ GMP-Mo, CD319^+^ MDP-Mo, and neutrophils were sorted to purity and activated by PMA for six hours. Staining for DNA, MPO, and citrullinated H3 revealed a significantly higher propensity of triple-positive events in GMP-Mo, compared to MDP-Mo (**Fig 3K**). This is in line with earlier reports of ET formation by human monocytes ^43^.

Finally, to compare their phagocytic potential, CD177^+^ and CD319^+^ BM monocytes were sorted to purity and incubated with GFP-expressing *Salmonella typhimurii* for two hours. Flow-cytometric analysis revealed that CD177^+^ GMP-Mo showed a superior capability to internalize bacteria (**Fig 3L, Suppl Fig 3M**). However, also MDP-Mo showed robust phagocytosis activity in this assay, in line with their monocytic nature.

Taken together, these *in vitro* assays suggest that, beyond their transcriptomic disparities, CD177^+^ GMP-Mo also display functional activities that differentiate them from MDP-Mo, including classical neutrophil activities, such as attraction by bacterial peptides, ET formation, and phagocytosis.

### GMP-Mo and MDP-Mo give rise to NCM and intestinal macrophages

A main homeostatic CM function is to maintain selected MΦ compartments, including cells in mucosal tissues ^1,44,45^. To investigate prospective fates of CD177^+^ GMP-Mo and CD319^+^ MDP-Mo, we isolated the two CM subsets based on surface marker expression from reporter animals that endow them with a discrete label, i.e. *Cx3cr1^Gfp^*, *Cx3cr1^Cre^:R26^LSL-TdTom^,* and *Ms4a3^Cre^:R26^LSL-TdTom^* mice ^11,13,46^, and performed adoptive transfer experiments. To free niches for engraftment ^47^, we took advantage of irradiation chimeras generated with *Cx3cr1^DTR^* BM. Specifically, diphtheria toxin (DTx) treatment of these animals results in depletion of monocytes and Cx_3_cr1^+^ MΦ populations ^48,49^.

Equal amounts of GMP-Mo and MDP-Mo (2 x 10^5^) were co-transferred into DTx-treated *Cx3cr1^DTR^* chimeras 8-10 weeks after irradiation and BM transfer (**Fig 4A**). Depletion of blood monocytes, including CM and NCM, by DTx was confirmed by flow cytometry (**Fig 4B**). Blood analysis of some recipient animals on day 1 after transfer, revealed circulating engrafted TdTom^+^ GMP-Mo and GFP^+^ MDP-Mo, at roughly equal numbers (**Fig 4C**). In line with earlier reports and indicating differentiation towards NCM ^18^, both cell types had begun to downregulate Ly6C (**Fig 4C**). When bled on day 3 after transfer, the analysis revealed that both GMP-Mo and MDP-Mo had converted into Ly6C^low^ NCM while maintaining the ratio (**Fig 4D**). Surface marker profiling revealed similar expression of the integrin CD11c among GMP-Mo and MDP-Mo derived NCM (**Fig 5E**). However, MDP-Mo NCM displayed significantly higher expression of PDL1, a recently reported NCM marker ^50^ (**Fig 4E**). Conversely, the classical monocyte markers CD115 and CD11b, were higher expressed on GMP-Mo derived NCM, albeit not reaching significance (**Fig 4E**).

**Figure 4:**
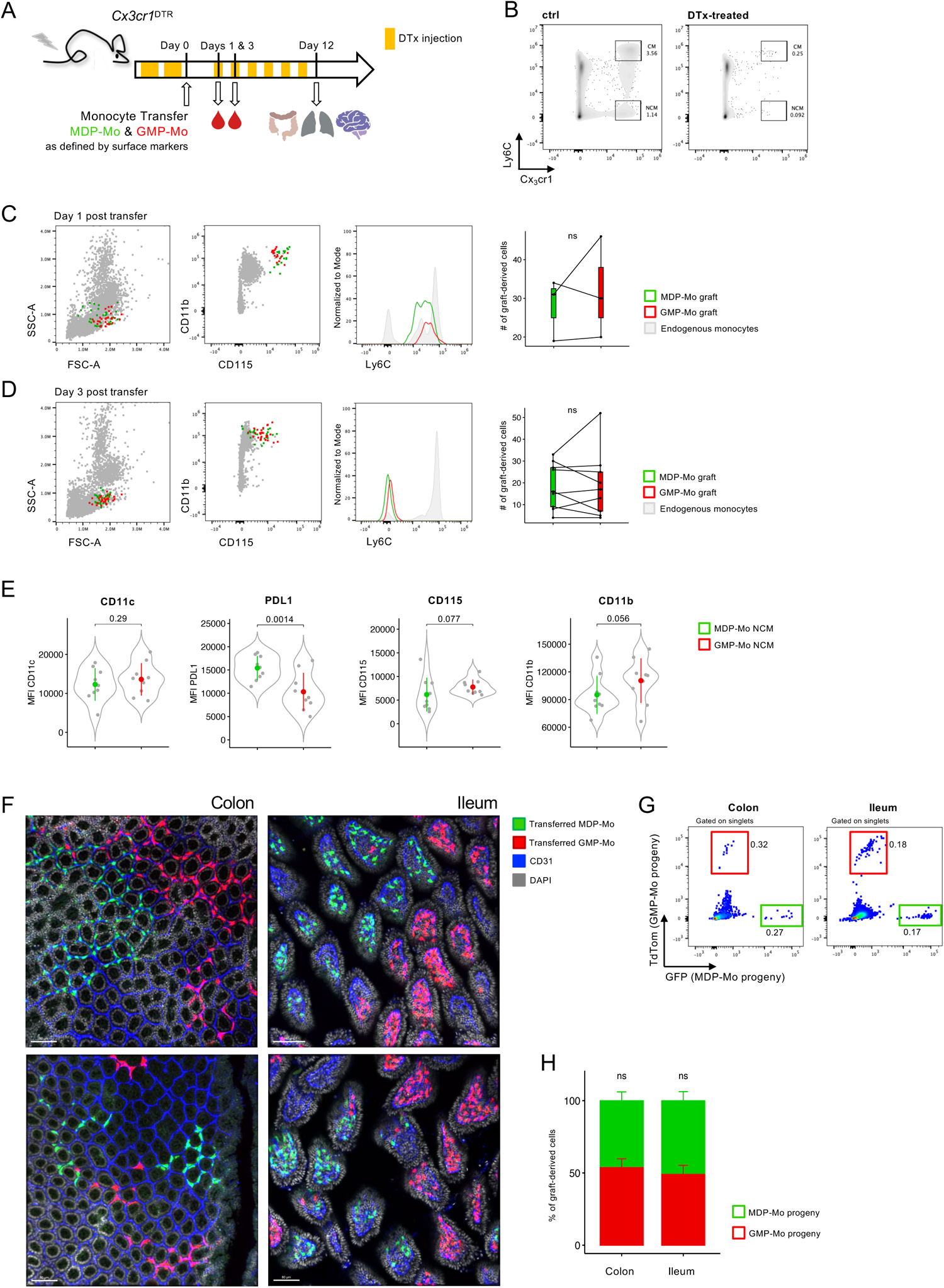
GMP-Mo and MDP-Mo give rise to NCM and intestinal macrophages. (A) Experimental scheme of adoptive transfer of 2×10^5^ GMP-Mo and MDP-Mo, each, as defined by surface markers. *Cx3cr1^DTR^*chimeras were depleted of Cx_3_cr1-expressing cells by repeated injection of 18 ng/g DTx / g bodyweight 8-10 weeks after chimerism. On day 0, CM subsets were adoptively co-transferred. Hereafter, mice were administered 12 ng/g DTx / g bodyweight every other day until sacrifice. (B) Blood analysis of DTx- and PBS-treated *Cx3cr1^DTR^* chimeras. (n=10 from 4 independent experiments) (C) Blood analysis of circulating graft cells one day after adoptive transfer. (n=3) (D) Blood analysis of circulating graft cells three days after adoptive transfer. (n=9) (E) Quantification of surface marker expression on graft cells in the blood three days after transfer. (n=9) (F) Microscopy images of graft-derived tissue-resident MΦ in the colon and ileum of recipient animals. Scale bar = 80-100 µm. (n=6 from 3 independent experiments) (G) Flow cytometry analysis of graft-derived tissue-resident MΦ in the colon and ileum of recipient animals. (n=4) (H) Quantification of the distribution of graft-derived tissue-resident MΦ in the colon and ileum of recipient animals. (n=7)

**Figure 5:**
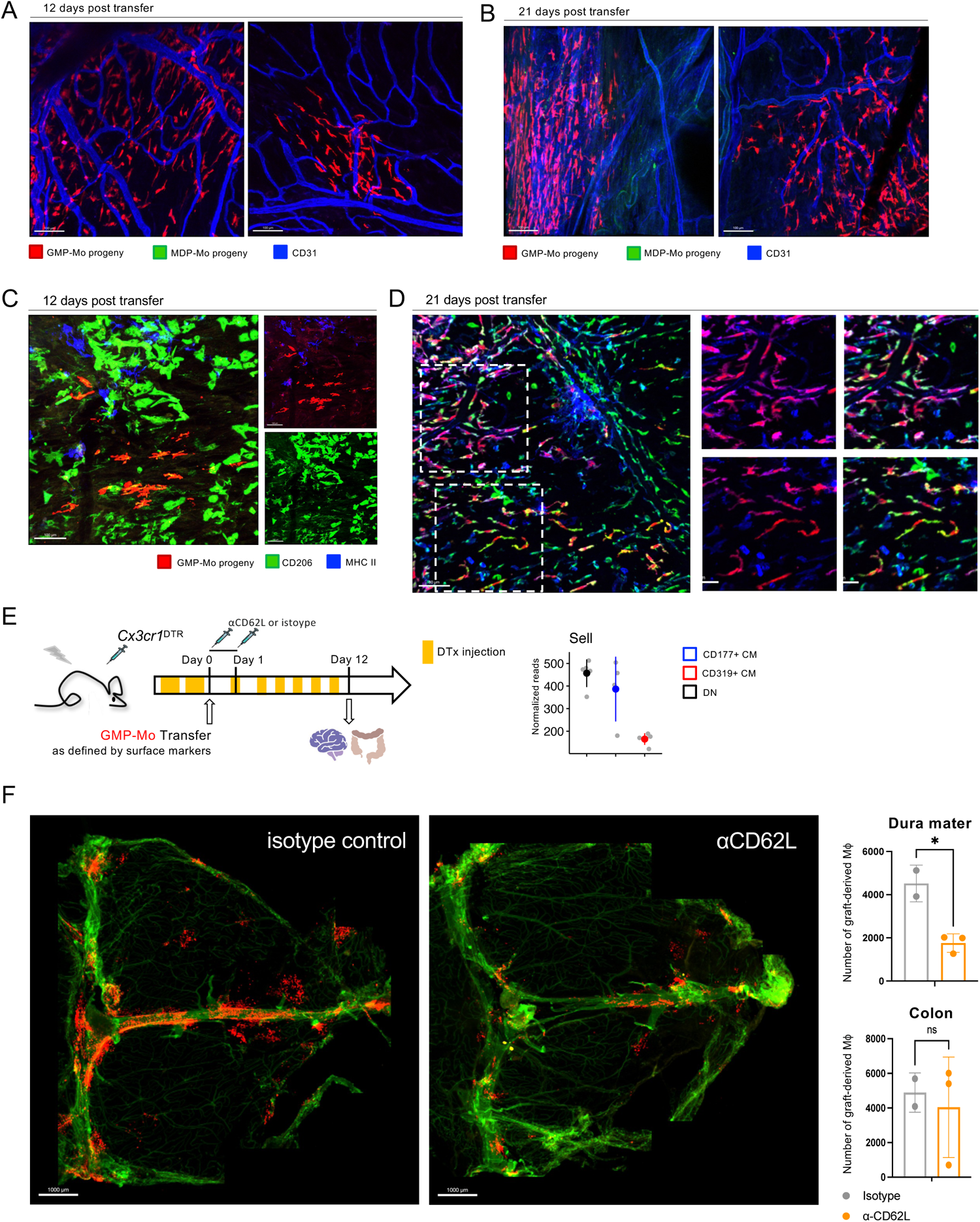
GMP-Mo but not MDP-Mo give rise to meningeal dura mater macrophages. (A) Microscopy image of the dura mater of recipient mice 12 days after transfer. Both images depict the peripheral dura. Scale bars = 100 µm. (n=12 from 4 independent experiments) (B) Microscopy image of the dura mater of recipient mice 21 days after transfer. Left: Sagittal sinus, right: periphery. Scale bars = 100 µm. (n=3) (C) Microscopy image of the dura mater. GMP-Mo-derived BAM were stained for CD206 (green) and MHC-II (blue), 12 days after transfer. Scale bar = 100 µm. (n=3) (D) Microscopy image of the dura mater. GMP-Mo-derived BAM were stained for CD206 (green) and MHC-II (blue), 21 days after transfer. Scale bar = 80 µm. (n=2) (E) Left: Schematic of the adoptive transfer experiment probing for the effect of CD62L on GMP-Mo recruitment to the dura mater. 2×10^5^ GMP-Mo were injected into each mouse. Mice were administered αCD62L or isotype antibody (100 µg i.v., each) 1 hour before and 24 hours after cell transfer. Right: Normalized reads for *Sell*, encoding CD62L among blood CM subsets, as assessed by bulk RNAseq (n=4-5). (F) Tile scans of the dura mater of αCD62L- and isotype-treated mice (left and middle) as well as quantification off total cell numbers detected in dura mater and colon in each treatment group (right). Scale bars = 1000 µm. (n=2-3)

On day 12 after transfer, several organs of the recipient mice were harvested to investigate the potential of GMP-Mo and MDP-Mo to give rise to long-lived tissue-resident MΦ ^18,47,51^. In both ileum and colon, clones of GMP-Mo and MDP-Mo derived MΦ were found and had clonally expanded, as reported earlier ^47^, (**Fig 4F**). Quantification of graft-derived MΦ by microscopy and flow cytometry revealed equal contributions of GMP-Mo and MDP-Mo to both ileal and colonic MΦ compartments (**Fig 4G,H**).

Collectively, we show that following engraftment, both GMP-Mo and MDP-Mo give rise to NCM and gut MΦ with comparable efficiency in a competitive setting. Of note, GMP-Mo and MDP-Mo derived NCM displayed distinct phenotypes, including PDL1 expression, suggesting that their derivation from GMP-Mo or MDP-Mo could have impact on NCM functions.

### GMP-Mo but not MDP-Mo give rise to meningeal dura mater macrophages

The MΦ compartment of the central nervous system (CNS) comprises parenchymal microglia, meningeal MΦ in dura mater and leptomeninges, as well as MΦ in the perivascular space and choroid plexus ^52,53^.

To probe for the ability of GMP-Mo and MDP-Mo to replenish CNS MΦ niches, we analyzed the brains of DTx-treated *Cx3cr1^DTR^* chimeras 12 and 21 days after engraftment (**Fig 4A**). The brain parenchyma of the recipient mice was devoid of graft-derived labelled cells, in line with DTx insensitivity of this compartment due to the limited replacement of radio-resistant microglia by DTR transgenic cells in the BM chimeras ^54^ (**Suppl Fig 4A**). In contrast, meningeal MΦ of the *Cx3cr1^DTR^* chimeras were partially depleted by the DTx regimen (**Suppl Fig 4B**). Nevertheless, no labelled cells were detected in the leptomeninges of the recipient mice (**Suppl Fig 4B**). Surprisingly, however, we observed an efficient repopulation of MΦ in the dura mater, albeit exclusively by TdTom^+^ GMP-Mo (**Fig 5A,B**). Specifically, cells concentrated along the sagittal sinus, but could also be found in more distant areas (**Fig 5A,B, Suppl Fig 4C**). Engraftment was sustained, since TdTom^+^ cells were still abundantly present 3 weeks after transfer, indicative of *in situ* proliferation of the graft (**Fig 5B**). Indeed, co-transfer of differentially labeled GMP-Mo revealed peripheral clusters of GFP^+^ or TdTom^+^ in the brains of recipient mice, suggesting clonal proliferation, as reported earlier for the gut ^47^ (**Suppl Fig 4D**). The inability of MDP-Mo to seed the dura meter was confirmed in independent transfer experiments in which we switched the labels, engrafting TdTom^+^ MDP-Mo and GFP^+^ GMP-Mo (**Suppl Fig 4E**). By day 12, GMP-Mo derived cells had acquired low levels of CD206 expression, a hallmark of dura mater MΦ ^53^ (**Fig 5C**). However, surface MHC II expression and higher levels of CD206 were only detected at day 21, suggesting further *in situ* maturation of the cells (**Fig 5D**).

To investigate the underlying mechanism of the selective dura mater seeding of GMP-Mo, but not MDP-Mo, we turned to the list of differentially expressed trafficking molecules. Specifically, we noted high level of expression of *Sell*, encoding for CD62L, by GMP-Mo, as compared to MDP-Mo, both on the transcriptional and protein level (**Fig 5E**), and as also shown via CITE-seq for the BM CM subsets (**Fig 1D**). To probe for a functional relevance of CD62L expression for the dura mater seeding by CM, we transferred labelled GMP-Mo into recipient mice which were i.v. treated on two consecutive days with αCD62L antibody (100 μg / mouse) or isotype control (**Fig 5E**). Unimpaired seeding of the colon by grafted cells at 12 days excluded cytotoxicity of the treatment (**Fig 5F, Suppl Fig 4F**). Analysis of the dura mater of the recipients after transfer revealed that αCD62L blockade led to a decrease of TdTom^+^ dura mater MΦ, compared to control mice (**Fig 4F**).

In conclusion, our data establish differential homing potential of the two CM subpopulations. Specifically, GMP-Mo but not MDP-Mo were able to repopulate an experimentally depleted dura mater MΦ niche, probably related to their expression of CD62L.

### GMP-Mo and MDP-Mo give rise to distinct macrophage populations in the lung

Pulmonary interstitial MΦ (IM) are heterogenous and comprise CD206^−^ cells and CD206^+^ IM ^55^, as well as lung-resident CD16.2^+^ cells related to NCM ^23^. The latter were considered IM precursors ^23^, a finding supported by observations from humanized animals ^24^, although more recent data suggest that IM develop from CM through a proliferating monocytic intermediate ^56^.

Engrafted TdTom^+^ GMP-Mo and GFP^+^ MDP-Mo derived cells could be readily detected in the recipient lungs by histology and flow cytometric analysis on day 12 (**Fig 6A, B, Suppl Fig 5A**), a time point when only rare grafted cells were discernable in the blood (**Suppl Fig 5B**). Interestingly, TdTom^+^ and GFP^+^ cells significantly differed with respect to their relative contribution to pulmonary IM subpopulations. Specifically, while both GMP-Mo and MDP-Mo efficiently reconstituted CD16.2^+^ cells, the majority of lung IM originated from MDP-Mo (**Fig 6C,D**). The less abundant GMP-Mo derived IM (GMP-IM) further differed from MDP-Mo derived IM (MDP-IM) by higher CD16.2 and lower MHC-II surface expression (**Fig 6E, Suppl Fig 5C**).

**Figure 6:**
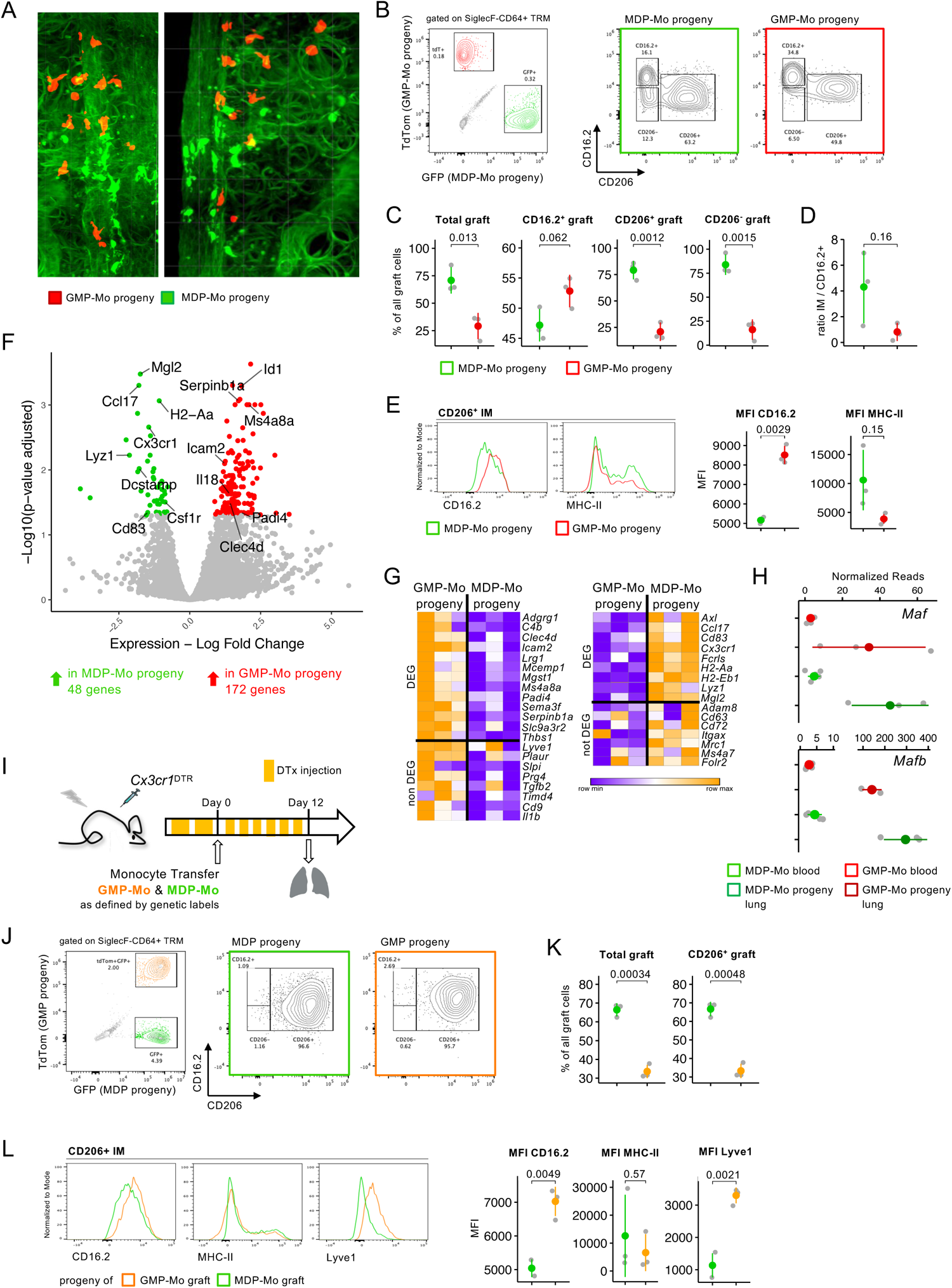
MDP-Mo preferentially give rise to lung interstitial macrophages. (A) Representative microscopy image of CUBIC-cleared lungs showing graft-derived cells in the parenchyma. (B) Flow cytometry plots of graft-derived cells in the lungs of recipient animals, and CD16.2 vs CD206 plots for each graft-derived population (n=6 from 2 independent experiments) (C) Quantification of the contribution of MDP-Mo and GMP-Mo derived populations to selected lung phagocyte populations, normalized to total number of graft-derived cells in the respective gates. (n=6 from 2 independent experiments) (D) Ratios of mature IM to CD16.2^+^ precursors for MDP-Mo- and GMP-Mo-derived graft cells. (n=6 from 2 independent experiments) (E) Histogram plots and quantification of the surface expression of CD16.2 and MHC-II on CD206^+^ MDP-Mo- and GMP-Mo-derived IM. (n=6 from 2 independent experiments) (F) Volcano plot of differentially expressed genes among total MDP-Mo and GMP-Mo derived phagocyte populations from the lung of recipient animals. (n=3) (G) Heatmap of selected differentially and non-differentially expressed genes attributed to Lyve1^+^MHC-II^low^ and Lyve1^−^MHC-II^hi^ IM as defined by Chakarov *et al.* (n=3) (H) Gene expression plots of *Maf* and *Mafb* among MDP-Mo- and GMP-Mo-derived IM and blood precursors. (n=3-5) (I) Experimental scheme for adoptive transfer of GMP- and MDP-derived monocytes from *Ms4a3^Cre^:R26^LSL-TdTom^:Cx3cr1^Gfp^* mice as defined by genetic labels. *Cx3cr1^DTR^* chimeras were depleted off Cx_3_cr1-expressing cells by twice injection of 18 ng/g DTx / g bodyweight 8-10 weeks after chimerism. On day 0, CM subsets were adoptively co-transferred. Hereafter, mice were administered 12 ng/g DTx / g bodyweight every other day until sacrifice. (J) Flow cytometry plots of graft-derived cells in the lungs of recipient animals, and CD16.2 vs CD206 plots for each graft-derived population. (n=6 from 2 independent experiments) (K) Quantification of the contribution of MDP- and GMP-derived populations to selected lung phagocyte populations, normalized to total number of graft-derived cells in the respective gates. (n=6 from 2 independent experiments) (L) Histogram plots and quantification of the surface expression of CD16.2, MHC-II, and Lyve1 on CD206^+^ MDP- and GMP-derived IM. (n=6 from 2 independent experiments)

To further characterize the pulmonary GMP-Mo and MDP-Mo progeny, we performed bulk RNAseq on engrafted TdTom^+^ and GFP^+^ cell retrieved from recipient lungs (**Suppl Fig 5D**). Out of a total number of 220 DEG, 48 genes were preferentially expressed in MDP-IM, while 172 genes displayed higher transcription in GMP-IM (**Fig 6F, Suppl Fig 5E**). Metascape analysis ^57^ revealed enrichment in *angiogenesis* and *wound healing* pathways in the IM progeny of GMP-Mo (**Suppl Fig 5F**). Conversely, profiles of MDP-IM displayed gene modules regulating the *immune response* and *leukocyte differentiation* (**Suppl Fig 5F**). Interestingly, the transcriptome of the GMP-Mo progeny showed significant overlap with an expression signature reported for Lyve1^+^MHC-II^low^ lung IM, while MDP-Mo progeny was more akin to Lyve1^−^MHC-II^hi^ lung IM ^58^ (**Fig 6G, Suppl Fig 5G,H**). Moreover, compared to GMP-IM, cells derived from MDP-Mo displayed twice higher expression of *Mafb*, a TF recently suggested to be required for CM differentiation into lung IM ^56^ (**Fig 6H**). Expression of c-Maf, encoded by *Maf* and proposed to imprint CD206^+^ IM identity ^56^, showed similar expression in IM derived from the two CM subsets (**Fig 6H**).

To provide further evidence for a link of CM ontogeny to the IM dichotomy ^58^, we next performed a competitive adoptive transfer of GMP- and MDP-derived monocytes isolated from BM of *Ms4a3^Cre^:R26^LSL-TdTom^:Cx3cr1^Gfp^*mice according to reporter expression (**Fig 6I**). MDP-derived GFP^+^ monocytes gave also in this setting preferentially rise to IM over GMP-derived TdTom^+^ GFP^+^ CM (**Fig 6J,K**). Furthermore, CD16.2 expression was higher on GMP-derived CD206^+^ IM, in agreement with our earlier observation (**Fig 6L**). Interestingly, and in line with the transcriptome data, GMP-Mo derived IM displayed higher surface expression of Lyve1. However, we observed no differential MHC-II expression among the graft-derived IM populations (**Fig 6L**).

In conclusion, using a competitive adoptive transfer approach, we establish that GMP-Mo and MDP-Mo have distinct differentiation trajectories upon their seeding of the lung, suggesting that CM fates in tissues are determined by ontogeny.

### GMP-Mo and MDP-Mo differentiation in steady-state lungs

Our adoptive transfer strategy and the associated conditioning bear inherent caveats. For once, irradiation induces low-grade inflammation in exposed organs ^59,60^, as does likely the DTx-induced cell ablation. Secondly, the transfer of equal numbers of GMP-Mo and MDP-Mo does not represent steady-state conditions. To circumvent these limitations, we adopted an irradiation strategy involving thorax shielding ^56,61^ and *Ms4a3^Cre^:R26^LSL-TdTom^:Cx3cr1^Gfp^* BM transfer (**Fig 7A, Suppl Fig 6A**), which yielded about 50% chimerism by week 4 (**Fig 7B, Suppl Fig 6B**). As in *Ms4a3^Cre^:R26^LSL-TdTom^:Cx3cr1^Gfp^*mice (**Fig 2A**), GFP^+^ MDP-Mo were rare in the circulation of the chimeras and vastly outnumbered by GFP^+^ TdTom^+^ GMP-Mo (**Fig 7C**). Unlike in the former mice, however, the reporter labels allow in the chimeras to unequivocally identify monocyte-derived cells and discriminate them from embryonic-derived tissue-resident MΦ.

**Figure 7:**
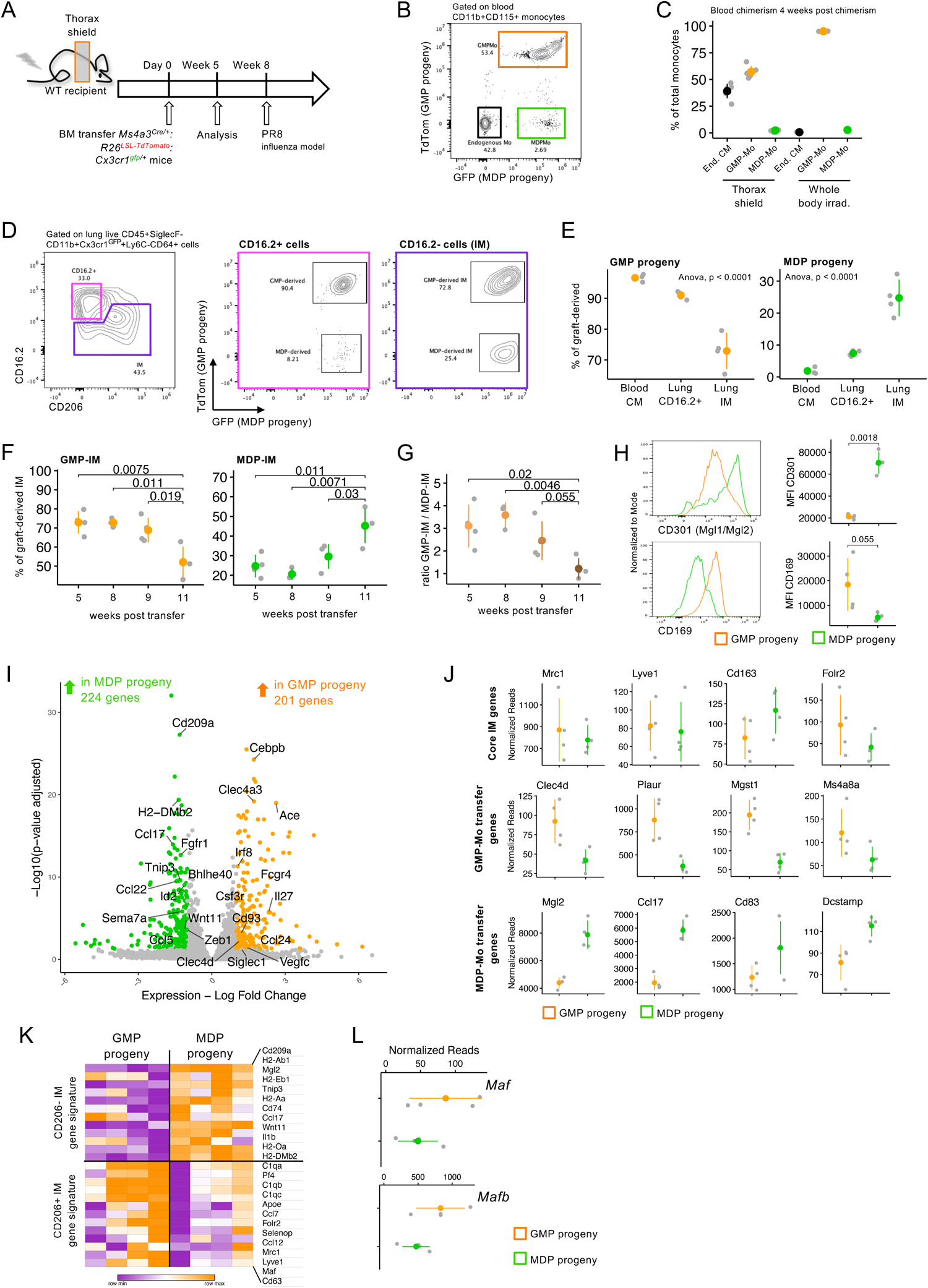
GMP- and MDP-Mo differentiation in steady-state lungs. (A) Experimental scheme of generation of thorax-shielded chimeras to study contributions of GMP-Mo and MDP-Mo to lung IM in steady-state and upon viral challenge. (B) Representative flow cytometry plot of blood endogenous and graft-derived monocytes in thorax-shielded recipients, 4 weeks following chimerism. (n=4 from 2 independent experiments) (C) Distribution of endogenous and graft-derived monocytes in thorax-shielded and whole-body-irradiated recipients, 4 weeks following chimerism. (n=2-4 from 3 independent experiments) (D) Representative flow cytometry plot of graft-derived CD64^+^ MΦ in thorax-shielded lungs (left), and the distributions of GMP- and MDP-derived cells in the CD16.2^+^ (middle) and CD16.2^−^ gates (right), 5 weeks following chimerism. (n=12 from 2 independent experiments) (E) Distribution of GMP- and MDP-derived cells among monocyte/ MΦ populations in the blood and lungs of thorax-shielded chimeras, 5 weeks following chimerism. (n=12 from 2 independent experiments) (F) Distribution of engrafted GMP- and MDP-derived IM in thorax-shielded recipients on different timepoints following chimerism. (n=20 from 2 independent experiments) (G) Ratio of GMP-IM and MDP-IM in thorax-shielded recipients on different timepoints following chimerism. (n=20 from 2 independent experiments) (H) Representative flow cytometry histograms and statistical analysis of selected surface markers on GMP-IM (orange) and MDP-IM (green). (n=4 from 2 independent experiments) (I) Volcano plot of differentially expressed genes among GMP-Mo and MDP-Mo derived CD64^+^ cells. (n=4) (J) Scatter plots of selected genes expressed in GMP- and MDP-derived CD64^+^ cells. *Clec4d, Plaur, Mgst1,* and *Ccl17* were differentially expressed. (n=4) (K) Heat map of CD206- and CD206+ IM gene signatures, as defined by Vanneste *et al.*, in GMP- and MDP-derived CD64^+^ cells. (n=4) (L) Scatter plots of *Maf* and *Mafb* expression in GMP- and MDP-derived CD64^+^ cells. (n=4)

Analysis of the graft-derived CD64^+^ lung cell compartment of the thorax-shielded chimeras 5 weeks after irradiation revealed a low contribution of engrafted MDP-Mo to CD16.2^+^ cells (**Fig 7D, Suppl. Fig 6C**), in line with the notion that these cells are related to blood NCM ^23^. In stark contrast, however, and defying their low abundance in the blood (**Fig 7C**), engrafted MDP-Mo substantially contributed to the CD16.2^−^ IM population, constituting an average of 25 % of the cells (**Fig 7D,E**).

Analysis of the thorax-shielded mice at different timepoints following irradiation revealed that contribution of MDP-Mo to lung IM further increased significantly with time (**Fig 7F,G**). Moreover, CD206 expression was similar among graft-derived IM subsets and unaltered with time (**Suppl Fig 6D**). MDP-Mo progeny among CD16.2^+^ cells remained notably low (**Suppl Fig 6E**). Comparative surface profiling of GFP^+^ TdTom^+^ and GFP^+^ IM, i.e. GMP- and MDP-derived MΦ, revealed higher expression of CD301 (Mgl1/Mgl2) (**Fig 7H**), in line with the earlier observed transcriptomic prevalence of *Mgl2* in MDP-IM (**Fig 6F**). Furthermore, GMP-IM stood out by higher expression of CD169 (**Fig 7H**). Of note, the phenotype of GMP- and MDP-derived IM thus showed overlap with the profiles of three previously reported IM populations^55^, with MDP-IM akin to a CD11c^+^ CD206^int/low^ IM1 population (**Suppl Fig 6F**).

Next, we sorted CD11b^+^ Ly6C^−^ CD64^+^ CD16.2^−^ graft-derived lung IM to purity and subjected them to bulk RNAseq (**Suppl Fig 6G**). 425 genes were found to be differentially expressed among GMP- and MDP-derived cells (**Fig 7I**). Both subsets equally expressed core IM markers, such as *Mrc1* (encoding CD206) and *Lyve1* (**Fig 7J**), in line with flow cytometry data (**Suppl Fig 6F**). Importantly, we also observed overlap with gene signatures of IM derived from adoptively transferred GMP-Mo and MDP-Mo, such as differential *Clec4d* and *Mgl2* expression (**Fig 7J**). Further, MDP-derived but not GMP-derived cells were enriched in a CD206^−^ IM gene signature, as defined before ^56^ (**Fig 7K**), potentially related to lower expression of *Mafb* and *Maf*, previously shown to imprint CD206^+^ IM identity ^56^ (**Fig 7L**). Of note, BM transfers and monocyte transfers yield different engraftment (**Suppl Fig 6H**). Accordingly, and despite enrichment for CD64^+^ cells, the bulk transcriptome GFP^+^ MDP progeny in the chimeras comprised transcripts related to MDP-derived DC, such as *Ccr7* and *Flt3* (**Suppl Fig 6I**). Lastly, the earlier transcriptomic alignment with the data reported by Chakarov et al. ^23^ (**Fig 6G**) and an enrichment for the ‘*regulation of neuron differentiation’* pathway in MDP-IM (**Suppl Fig 6J**) prompted us to trace the distances of GMP- and MDP-derived cells to neurons. Yet, unbiased distance analysis of lung sections of the thorax-shielded chimeras did not yield evidence for differential anatomic locations of TdTom^+^ GFP^+^ GMP-IM and GFP^+^ MDP-IM with respect to Tubb3^+^ neurons (**Suppl Fig 6K**).

In conclusion, we establish that, although MDP-Mo represent a minor monocyte population in the circulation, they harbor a profound potential to contribute to the steady-state turnover of lung-resident IM. Moreover, the combined results of the adoptive transfers and the chimera analysis show that GMP and MDP-derived pulmonary MΦ significantly differ with respect to phenotype and transcriptional landscapes.

### GMP-Mo and MDP-Mo differentiation in challenged lungs

The above data corroborate the differential contributions of GMP- and MDP-derived CM to the healthy pulmonary IM compartment and establish that MDP-Mo, despite their sparsity in the blood, significantly participate in the homeostatic turnover of specific MΦ.

To investigate GMP-Mo and MDP-Mo contributions following challenge, we infected thorax-shielded [*Ms4a3^Cre^:R26^LSL-TdTom^:Cx3cr1^Gfp^*> WT] chimeras with a murine influenza virus (PR8 strain) (**Fig 8A**). The ratio of GMP-Mo and MDP-Mo in the circulation was unaltered by the viral challenge on the day of peak infection, as assessed by weight loss (**Suppl Fig 7A**), although MDP-Mo were elevated at d23 (**Suppl Fig 7B,C**). However, MDP-derived IM had proportionally decreased compared to control, likely due the high abundance of infiltrating GMP-Mo (**Fig 8B, Suppl Fig 7D**). This alteration was however transient, as following resolution of the inflammation, MDP-IM had recovered, with the GMP-IM / MDP-IM ratio on d23 following viral challenge being similar to that of d8 PBS-treated mice (**Fig 8C**).

**Figure 8:**
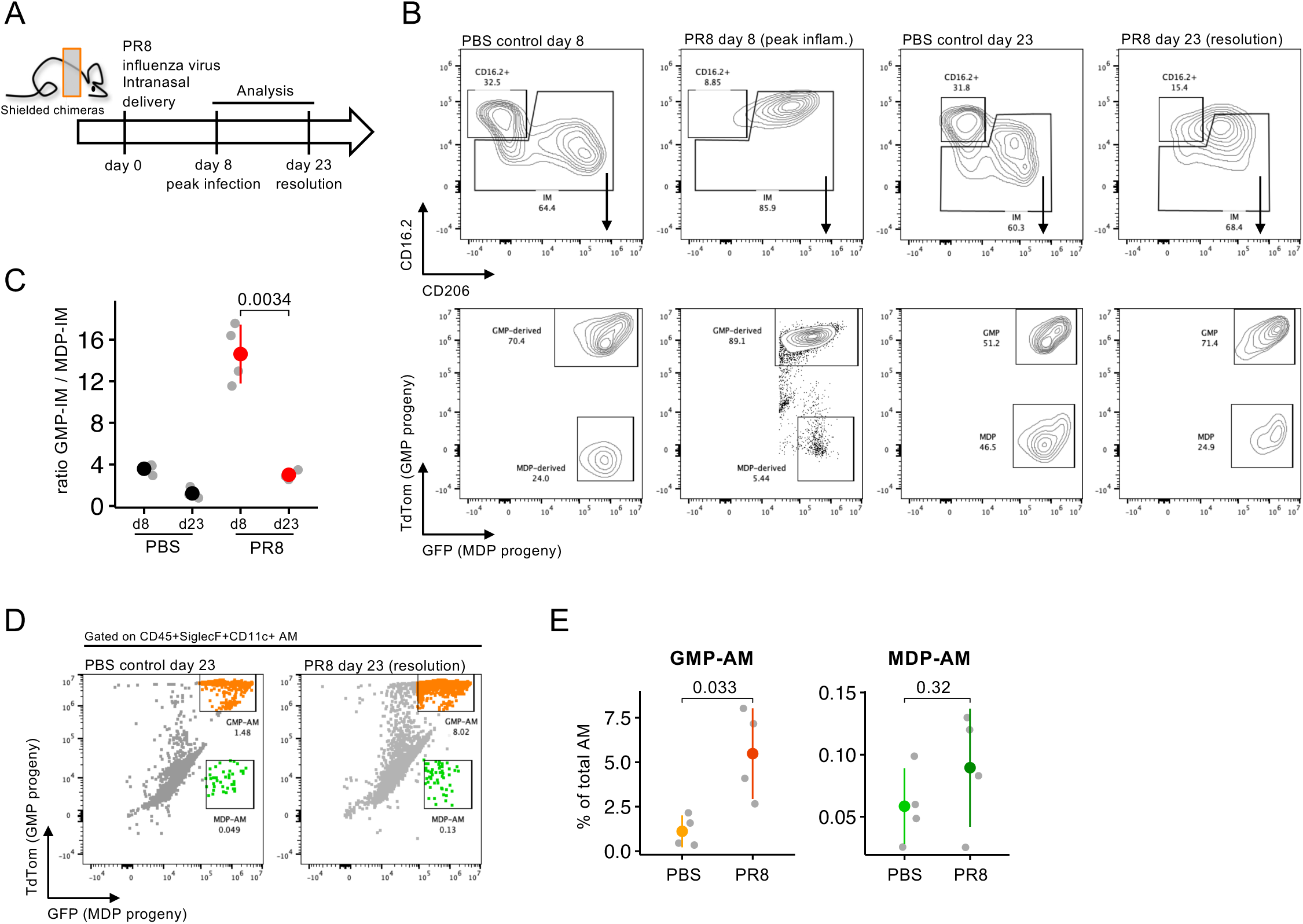
GMP-Mo and MDP-Mo differentiation in challenged lungs. (A) Experimental scheme of the influenza infection (PR8 strain) in thorax-shielded chimeras. (B) Representative flow cytometry plots of total graft-derived lung MΦ (upper row) and the distribution of GMP- and MDP-derived IM (lower row) in control (PBS) and challenged (PR8) thorax-shielded chimeras on different timepoints. (n=3-4 per condition) (C) Ratio of GMP- and MDP-derived IM across different conditions and timepoints in control (PBS) and challenged (PR8) thorax-shielded chimeras. (n=3-4 per condition) (D) Representative flow cytometry plots of AM in the lungs of thorax-shielded chimeras infected with PR8 or control (PBS) 23 days post challenge. (n=4 per condition) (E) Statistical analysis of the frequency of GMP- and MDP-derived AM among total AM in the lungs of thorax-shielded chimeras infected with PR8 or control (PBS) 23 days post challenge. (n=4 per condition)

Alveolar MΦ (AM) in unchallenged thorax-shielded chimeras were largely of host origin, in line with their fetal liver derivation ^62^, with rare adult monocyte descendants being GMP-Mo derived (**Fig 8D,E**). In contrast, and as reported ^63^, influenza-infected mice showed a prominent acute monocyte infiltration. Of note, also this replacement of AM following influenza challenge was predominantly mediated by GMP-Mo (**Fig 8D,E**).

In summary, we demonstrate that GMP-Mo are the dominant IM and AM precursor in influenza-challenged lungs. However, MDP-Mo significantly contributed to the lung IM compartment following resolution of inflammation.

## Discussion

Here, we extend recent pioneering reports on CM heterogeneity ^26,27^ and we establish shared and differential *in vivo* fates of GMP-and MDP-derived monocytes in selected tissues using adoptive transfer experiments, as well as thorax-shielded chimeras mimicking a homeostatic lung environment.

Monocyte subsets were first identified in human blood ^5^, and the subsequent definition of their murine correlates in *Cx3cr1^Gfp^* reporter mice ^6,7,46^ paved the way for functional studies of these cells in organismal context. Specifically, Geissmann and colleagues showed that murine Ly6C^low^ CCR2^−^ monocytes monitor blood vessel walls ^9^. Ly6C^high^ CCR2^+^ ‘inflammatory’ monocytes, on the other hand, perform classical monocyte functions, including, alongside neutrophils, the recruitment to sites of acute inflammation to promote and resolve inflammation. CM also give rise to long-lived MΦ at sites of injury ^4^ and, even in absence of overt inflammation, progressively replenish selected tissue MΦ compartments replacing embryo-derived cells ^47,64^. Exchange of YS-derived tissue MΦ could have long-term impact on the physiological state of tissues during aging. Moreover, unlike YS-derived MΦ, CM are, as HSC progeny, targets of somatic mutations associated with age-related clonal hematopoiesis (CH). Indeed, emerging evidence indicates that CH-afflicted monocyte-derived MΦ (MoMΦ) can contribute to cardiovascular and CNS pathologies ^65–67^.

Recent studies revealed transcriptomics-based evidence for CM heterogeneity with neutrophil and DC-like signatures as a function of distinct developmental pathways ^26,27^. We report surface markers that discriminate these proposed GMP-Mo and MDP-Mo among murine Ly6C^high^ BM and blood CM. We show that, in steady state, the vast majority of murine CM display a GMP-Mo signature, at least in mice kept under hyper-hygienic special pathogen free (SPF) conditions. GMP-Mo / MDP-Mo ratios might however differ in outbred animals roaming in the wild, given that the relative abundance of the subsets is influenced by environmental factors ^26^.

We define the SLAM family member 7 (CD319) as a marker for MDP-Mo. Notably, however, the discrimination of MDP-Mo and DC, including their precursors, remains a formidable challenge as these cell types share many surface markers, likely as a result of their common MDP ancestry ^2,26,30^. In our study, the definition of CM as CD11c^−^ cells ^10^ removed a considerable contamination of Ly6C^+^ Zbtb46^+^ Flt3^+^ cells, presumably cDC2 precursors, from the blood CM gate. Accordingly, the DC lineage-defining Zbtb46 TF was undetectable by deep sequencing of Ly6C^high^ CD319^+^ CD11c^−^ MDP-Mo. Moreover, we demonstrate that these cells are MΦ precursors and thus display a unique monocyte feature. For all the above we conclude that MDP-Mo, as defined in this study, are *bona fide* monocytes and not DC. We used the GPI-linked cell surface glycoprotein CD177 as a marker for GMP-Mo ^68^, although our adoptive precursor transfer experiments subsequently revealed that both GMP- and MDP-derived CM express comparable levels of CD177. Thus, CD177 alone cannot serve as a definitive GMP-Mo marker without additional discrimination by CD319.

Yáñez *et al.* demonstrated through adoptive cell transfers that CM can originate from both GMP and MDP ^26^. Supporting this notion, GMP-Mo were recently shown to derive from pro-neutrophils downstream of GMP in inflammatory conditions ^29^. Using *Ms4a3^Cre^:R26^LSL-TdTom^* mice, an established GMP fate mapping model ^13^, we could corroborate this developmental scheme. In line with the transcriptomes of CD177^+^ and CD177^−^ CD319^−^ GMP-Mo, more than 95% of blood CM were double labeled in the *Ms4a3^Cre^:R26^LSL-TdTom^:Cx3cr1^Gfp^*mice. This further substantiates that most CM in SPF-housed C57BL/6 animals constitute GMP-Mo. Conversely, a small fraction of GFP-only labeled CM of the reporter animals expressed higher levels of CD319 and Cx_3_cr1, suggesting that most MDP-Mo originate from a *Ms4a3*-independent progenitor, hence, the MDP ^26,27^. However, it should be noted that we found that GMP-derived CM also comprised a small fraction of CD319^+^ cells. It remains to be determined if this is due to reported minor activity of the *Ms4a3* gene in maturing CM originating from *Ms4a3^neg^* MDP ^13^.

Based on phenotypic and transcriptional features, the majority of CM in adult C57BL/6 mice kept under hyper-hygienic conditions consists of GMP-Mo. Following challenges, however, including CpG and IFNγ treatment, MDP-Mo become more dominant ^26^. A recent study has highlighted the importance of IFNγ in the differentiation of CM into inflammatory CNS-resident MoMΦ ^69^, in line with a report on pathogenic *Cxcl10^+^* CM emerging during autoimmune neuroinflammation ^70^. Of note, SLAMF7^+^ pro-inflammatory MΦ were shown to be IFNγ-dependent and dominant in human patients with rheumatoid arthritis, COVID-19, or inflammatory bowel disease ^71^. Finally, Goodridge and colleagues recently reported that MDP-Mo accumulate in aged mice, as assessed by transcriptional profiling and MHC-II surface expression ^72,73^. Clearly, MDP-Mo require further in-depth study, including their unequivocal delineation from cDC and their precursors.

The main homeostatic function of monocytes is arguably the differentiation into tissue-resident MΦ and our competitive adoptive cell transfer experiments establish that an equal proportion of GMP-Mo and MDP-Mo contribute to blood NCM and gut MΦ. However, it remains to be defined whether GMP-Mo and MDP-Mo progeny are identical, or differ such as we show with respect to NCM phenotypes. In line with this notion, we found GMP-Mo, but not MDP-Mo to give rise to dura mater MΦ, possibly in relation to differential CD62L expression. Differential adhesion molecule expression by GMP-Mo might also confer selective extravasation to other peripheral tissues.

We establish that MDP-Mo preferentially gave rise to lung IM in the competitive adoptive transfer, as well as a model of thorax-shielded chimeras in which labeled GMP-Mo and MDP-Mo circulate at steady-state ratios. Despite their scarcity in the blood, MDP-Mo gave rise to a substantial proportion of IM, steadily increasing with time. Pulmonary GMP-IM and MDP-IM also differed according to phenotypes and transcriptomes, that in part align with earlier reported expression profiles of lung IM populations ^56,58^. The absence of evidence for differential anatomic locations of GMP-IM and MDP-IM with respect to nerves in our study suggests a dominant impact of their CM origin as compared to instruction by niches. However, this aspect requires further investigation.

We show that MDP-Mo prominently contribute to the lung IM under homeostasis. In contrast, following an acute viral challenge it is GMP-Mo that are recruited and that differentiate into IM and AM. Interestingly though, MDP-IM proportions recovered upon resolution of inflammation, although it remains to be defined whether recruited GMP-IM are short-lived, or late MDP-Mo influx recovered the MDP-IM compartment.

What’s in a name? Combined with the earlier seminal work ^26,27^ we establish here that the murine CM compartment comprises two subsets that are derived from GMP and MDP and display distinct transcriptomic signatures, respectively. Goodridge and Klein accordingly referred to a neutrophil-like and DC-like dichotomy, and we show that GMP-Mo indeed display activities associated with neutrophils, such as ET formation and recruitment by fMLP, that could justify the term NeuMo. However, the DCMo term implies a function that currently lacks support by experimental evidence. We therefore refer to the cells throughout as GMP-Mo and MDP-Mo focusing on origin. Lastly, the segregation of the blood CM by CD177 and CD319 surface markers revealed a sizeable fraction of cells negative for both antigens. However, our profiling data suggest that CD177^+^ and ‘double negative’ CD177^−^ CD319^−^ CM are both GMP-Mo and hence might well have similar functions in homeostasis. Given their abundance and recruitment following challenge it is GMP-Mo that best fit the bill of ‘inflammatory monocytes’ ^26,27^, while the role of MDP-Mo in inflammation requires further study.

In conclusion, we have identified surface markers that delineate two previously reported murine CM subsets ^26,27^. We demonstrate that GMP-Mo and MDP-Mo are *bona fide* monocytes that recirculate in the blood and have the capacity to give rise to tissue resident MΦ. Surprisingly though, GMP-Mo and MDP-Mo show distinct homing and differentiation potential in peripheral tissues, such as the lung and meninges. As such, our data link CM dichotomy to tissue MΦ heterogeneity. This finding, together with the dynamic abundance of GMP-Mo and MDP-Mo following challenges, could have major impact on the long-term composition of tissue macrophages in given organs.

## Data deposit

Gene expression data sets are deposited at GEO (Accession # xxxx)

## Supplementary Data

**Suppl Table 1.** List of CITE-seq antibodies used in this study

## Author contributions

S. T. and S. J. conceived the project and designed experiments. S.T., J-S. K., N. L., A. R.,J. B., M. H., D. A., and M. G-V. performed experiments. S. T., J-S. K., and N. L. analyzed experiments. S. T. and A. S. performed the bulk RNAseq and bioinformatic analysis. J. B. performed the single-cell experiment. D. K., J. B., and K. M. analyzed the single-cell experiment. J-S. K. and N. L. performed imaging experiments. S. T. performed adoptive cell transfers. S. T. and S. J. wrote the manuscript. S. J. supervised the project.

## Acknowledgements

We would like to thank all members of the Jung laboratory, as well as the Jung lab alumni A. Mildner and S. Yona for advice and comments on the manuscript. We thank the staff of the Weizmann Animal and flow cytometry facilities. S.J is the incumbent of the Henry HG. Drake Professional Chair of Immunology. This research was generously supported by Morris Kahn Institute for Human Immunology. This project was funded by the Deutsche Forschungsgemeinschaft (DFG, German Research Foundation), Project-ID 259373024 – TRR 167, the Vienna Science and Technology Fund (WWTF), Project 713745, and the Israeli Science Foundation (ISF) (grant # 696/21).

## Supplementary Figure legends

**Supplemental Figure 1.**
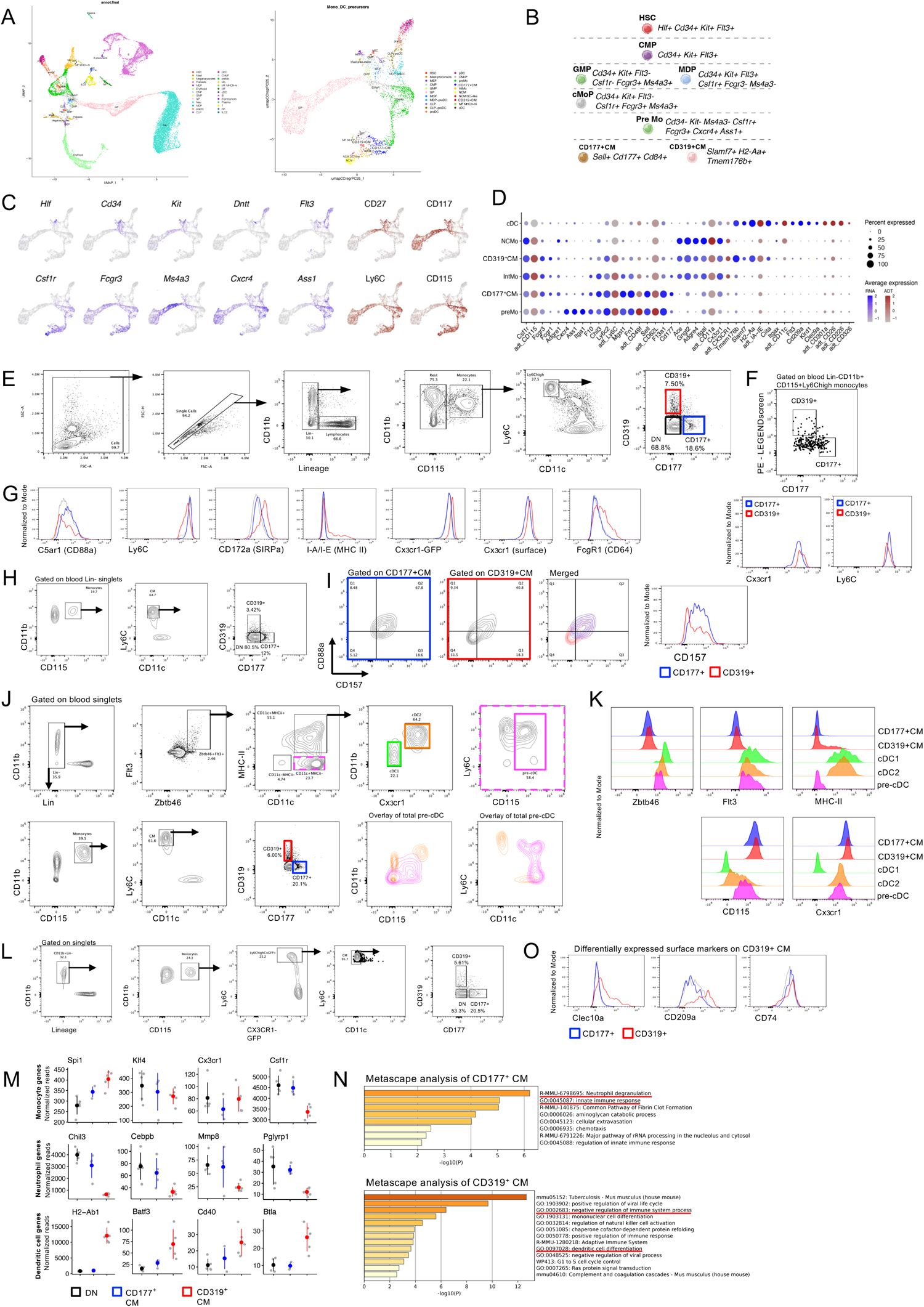
(A) UMAP plots of all identified BM cells. Monocytes, DCs and their precursors were selected and analyzed separately, as shown on the right. (B) Marker genes of BM cell types. (C) Expression of selected marker genes and proteins (the latter based on CITEseq antibodies) in the dataset from A. (D) Dot plot showing specific upregulated genes (blue) and proteins (red) for the individual monocyte and cDC subsets as identified by Cite-seq. Dot size represents the percentage of cells expressing the gene and colour represents its average expression. (E) Gating strategy for identification of blood immune subsets. (F) Dot plot from the LEGENDscreen identifying CD319 as blood CM marker. Histograms for Cx_3_cr1 and Ly6C shown as a validation for differences to CD177^+^ cells. (G) Distribution of selected markers for GMP-Mo and MDP-Mo on CM subsets as predicted by Weinreb *et al.* (n=3-5) (H) Gating strategy for CD177^+^ and CD319^+^ from the VIB mouse facility in Brussels, Belgium. (n=3) (I) Expression of CD88a and CD157 on GMP-Mo and MDP-Mo in the blood. (n=3) (J) Gating strategy for identification of a possible contamination of Zbtb46^+^Flt3^+^ cells in the blood CM gate used in this study. (n=3) (K) Histograms showing the distribution of DC and monocyte markers on monocyte and DC subsets in the blood. (n=3) (L) Gating strategy for sorting CM subsets from the blood of *Cx3cr1^Gfp/+^* mice for bulk RNAseq. (n=4-5) (M) Scatter plots of selected genes among CD177^+^ CM, CD319^+^ CM, and DN CM. Selected ‘monocyte genes’ were not differentially expressed among subsets. (n=4-5) (N) Metascape analysis of differentially expressed genes in CD177^+^ CM and CD319^+^ CM.

**Supplemental Figure 2.**
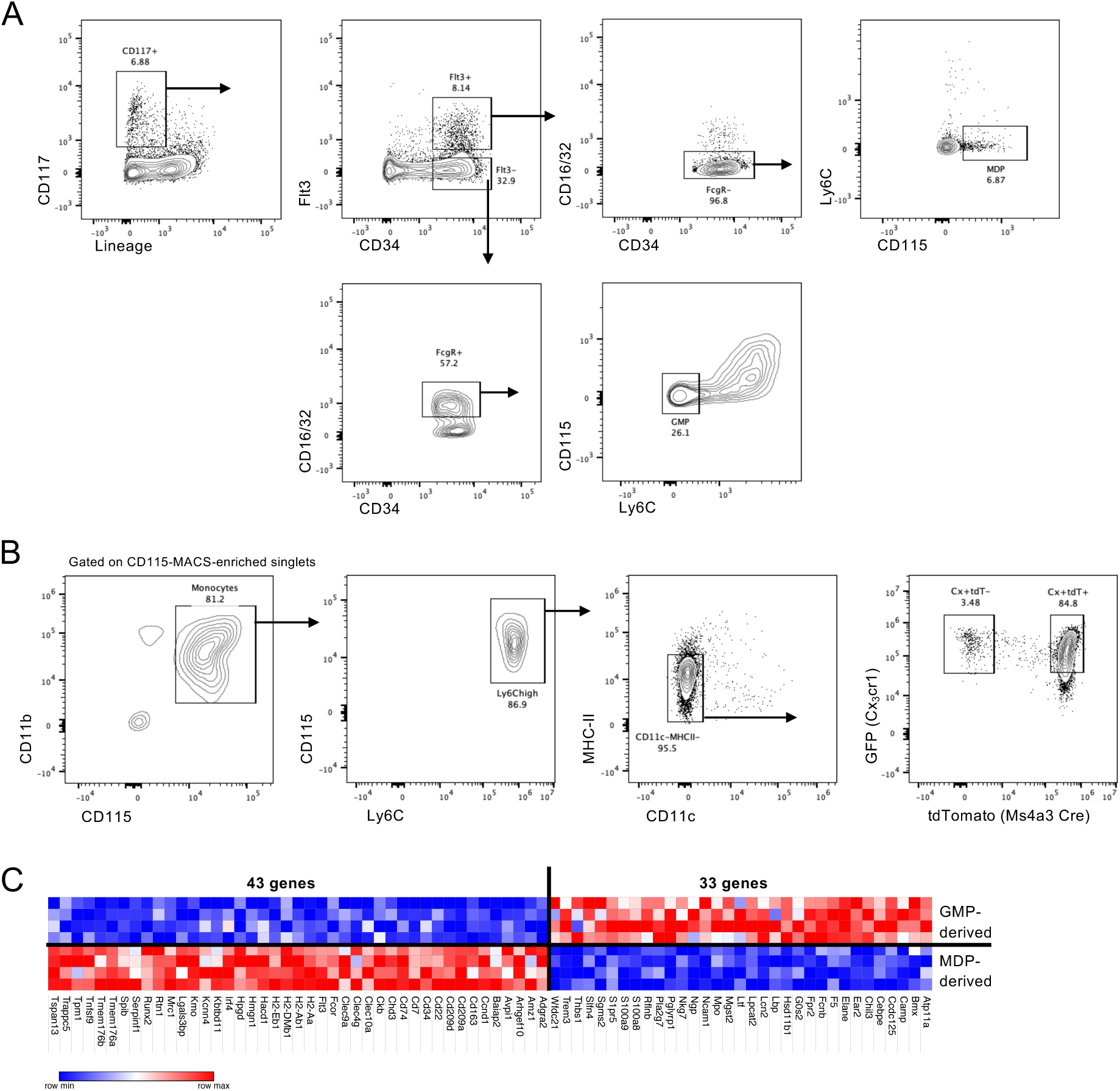
(A) Sorting strategy for adoptive transfer of GMP and MDP. (B) Gating strategy for sorting GMP- and MDP-derived CM from the BM of *Ms4a3^Cre^:R26^LSL-TdTom^:Cx3cr1^Gfp^* mice. (C) Heatmap of all DEG among GMP- and MDP-derived BM CM. (n=4)

**Supplementary Figure 3.**
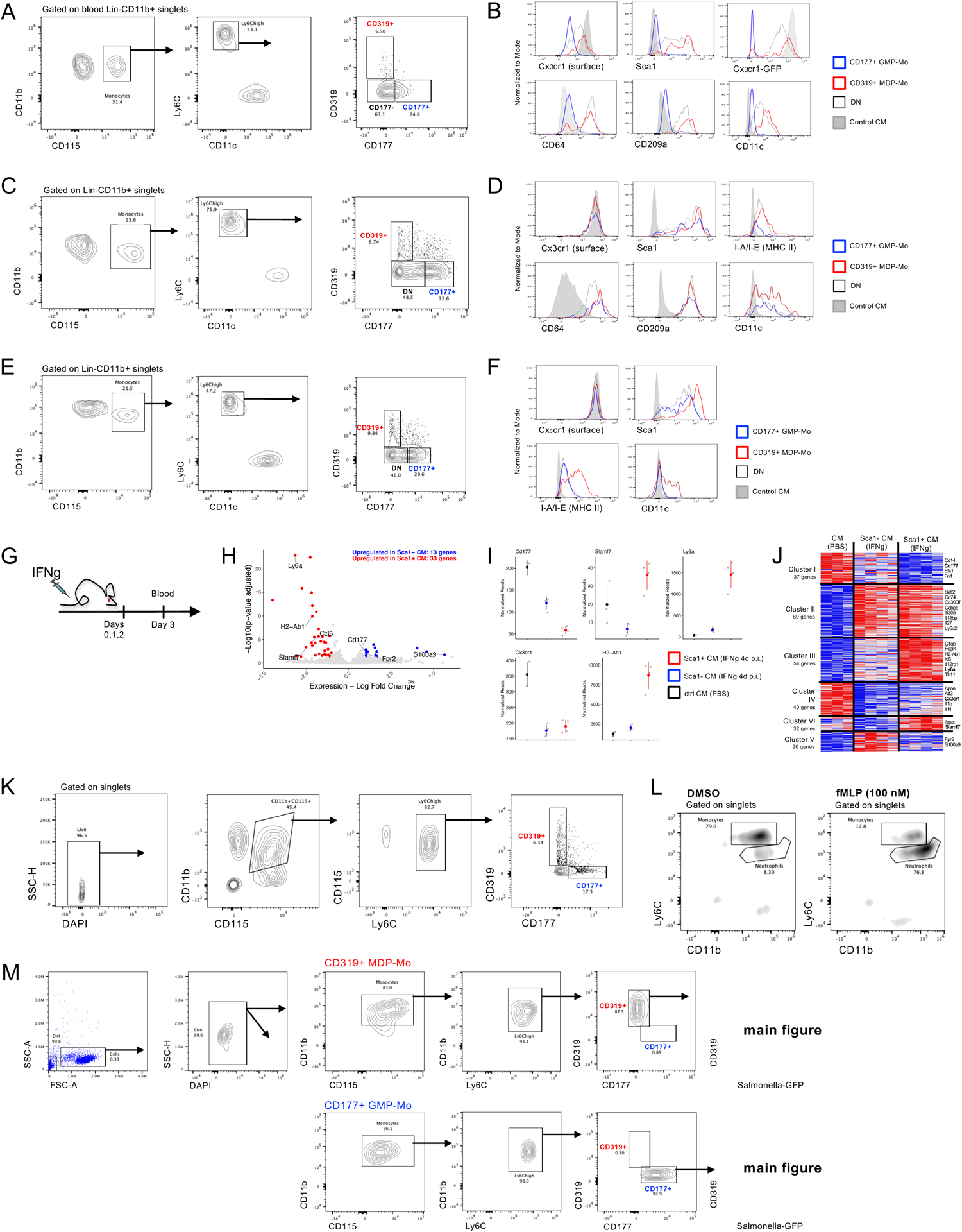
(A) Gating strategy for PBS-treated mice in the LPS experiment. (n=9 from 3 independent experiments) (B) Histograms for various markers on CM subsets in LPS-treated mice. (n=9 from 3 independent experiments) (C) Gating strategy for PBS-treated mice in the CpG experiment. (n=6 from 2 independent experiments) (D) Histograms for various markers on CM subsets in CpG-treated mice. (n=6 from 2 independent experiments) (E) Gating strategy for PBS-treated mice in the IFNγ experiment. (n=6 from 2 independent experiments) (F) Histograms for various markers on CM subsets in rIFNγ-treated mice. (n=6 from 2 independent experiments) (G) Experimental scheme for the sorting of Sca1^+^ and Sca1^−^ CM after rIFNγ treatment on 3 consecutive days. (H) Volcano plot of DEG genes among Sca1^+^ and Sca1^−^ CM from rIFNγ -treated mice. (I) Gene expression plots of selected genes from Sca1^+^ and Sca1^−^ CM from rIFNγ-treated mice. (J) Heat map of DEG among CM from control and rIFNγ -treated mice. (K) Gating strategy for sorting GMP-Mo and MDP-Mo from the BM for functional assays. (L) Density plots of migrated cells from DMSO (ctrl) and fMLP wells. (n=6 from 2 independent experiments) (M) Gating strategy for measuring engulfment of *S. Tm* in sorted GMP-Mo and MDP-Mo. (n=13 from 2 independent experiments)

**Supplementary Figure 4.**
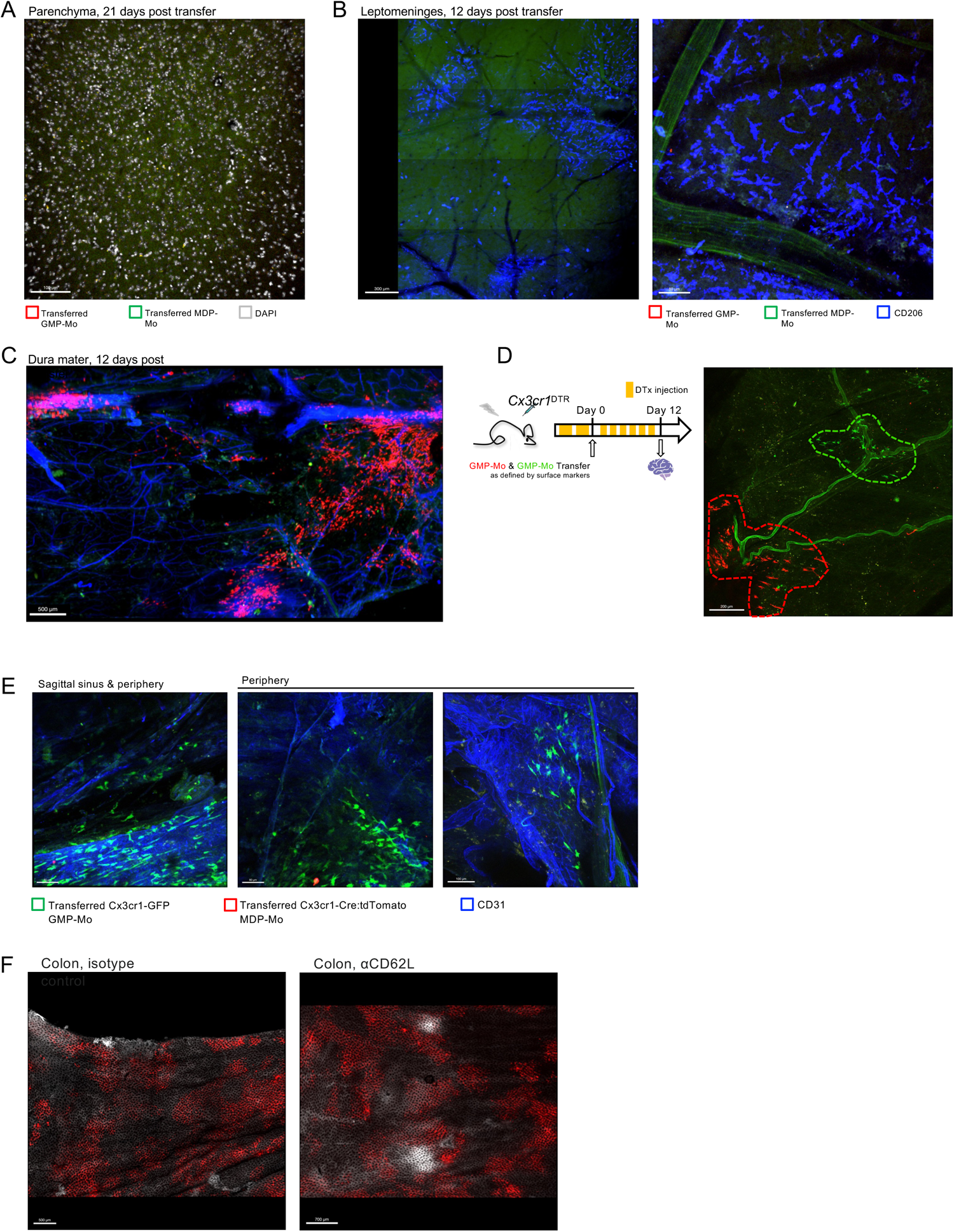
(A) Microscopy picture of the brain parenchyma of DTx-treated recipient mice. Scale bar = 100 µm. (n=3) (B) Microscopy picture of the leptomeninges of DTx-treated recipient mice. Scale bar = 300 µm. (n=3) (C) Tile scan of the dura mater of DTx-treated recipient mice. Scale bar = 500 µm. (n=1) (D) Experimental scheme and microscopy picture to show clonal expansion of GMP-Mo in the dura mater of macrophage-depleted recipients. Scale bar = 200 µm. (n=1) (E) Microscopy pictures of the dura mater of DTx-treated mice transferred with *Cx3cr1^Gfp^*GMP-Mo and *Cx3cr1^Cre^:R26^LSL-TdTom^* MDP-Mo (2×10^5^ each). Scale bars = 80-100 µm. (n=3) (F) Tile scans of the colons of isotype- and aCD62L-treated mice. Scale bars = 500-700 µm. (n=2-3)

**Supplementary Figure 5.**
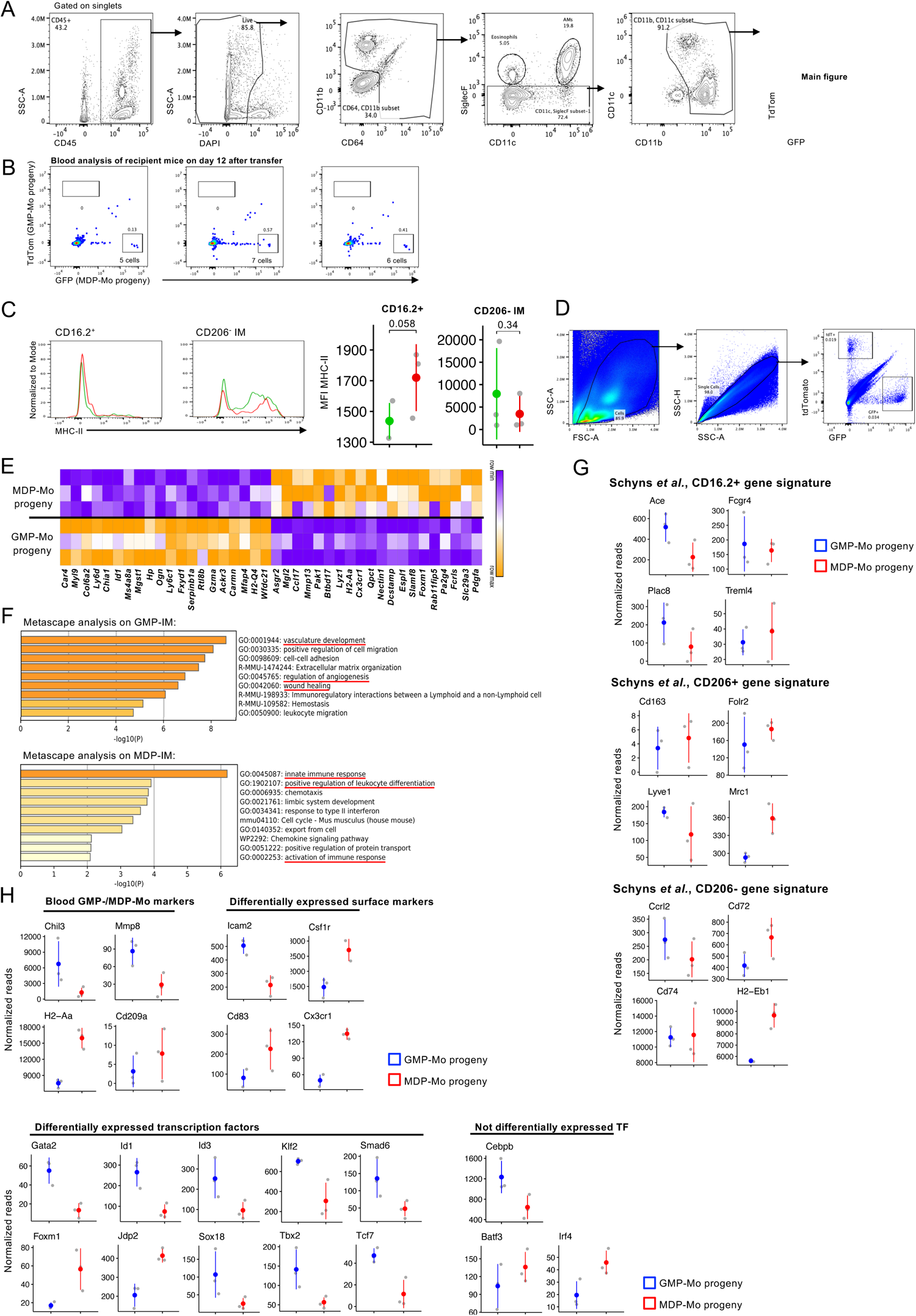
(A) Gating strategy for lung IM. (B) Blood of recipient mice on day 12 after transfer. Each plot represents a biological replicate. (C) Expression of MHC-II on the surface of CD16.2^+^ precursors and CD206^−^ IM, including MFI quantification. (n=6 from 2 independent experiments) (D) Sorting strategy for purification of GMP-Mo and MDP-Mo derived IM for bulk RNAseq analysis. (E) Heatmap of the 30 top DEG from GMP-IM and MDP-IM. (n=3) (F) Metascape analysis of DEG in GMP-IM and MDP-IM. (G) Gene plots for gene signatures of lung IM and precursors from Schyns *et al.* (n=3) (H) Gene plots for selected genes of interest among GMP-IM and MDP-IM. Among GMP-Mo / MDP-Mo markers’, *Mmp8* and *H2-Aa* were differentially expressed. (n=3)

**Supplementary Figure 6.**
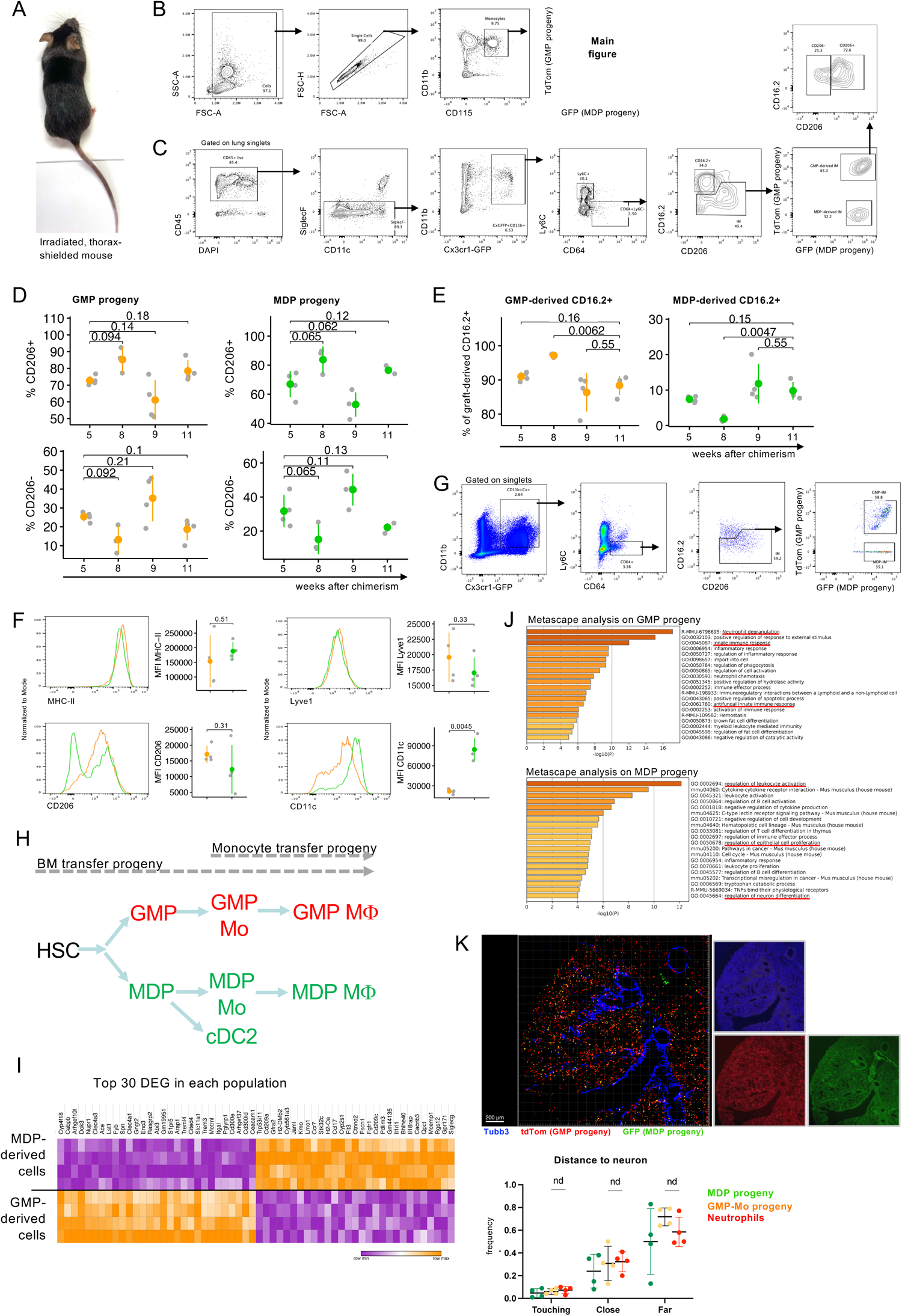
(A) Representative picture of an irradiated, thorax-shielded mouse, 8 weeks following irradiation. Note the black fur surrounding the thoracic cavity, indicating absence of irradiation in this area. (B) Gating strategy for assessing chimerism in blood monocytes. (C) Gating strategy for identification of graft-derived (GFP^+^) MΦ (Ly6C^−^CD64^+^) in recipient lungs. (D) Frequency of CD206^+^ and CD206^−^ GMP-IM and MDP-IM on different timepoints following chimerism. (n=20 from 2 independent experiments) (E) Distribution of engrafted GMP- and MDP-derived CD16.2^+^ cells in thorax-shielded recipients on different timepoints following chimerism. (n=20 from 2 independent experiments) (F) Expression of IM1,2,3 markers on GMP-IM and MDP-IM in thorax-shielded recipients. (n=4 from 2 independent experiments) (G) Sorting strategy for purification of GMP-Mo and MDP-Mo derived IM from thorax-shielded chimeras, later subjected to bulk RNAseq. (n=4) (H) Scheme of expected GMP and MDP progenies in the lungs of the adoptive transfer model (Figure 6) and the thorax-shielded BM chimeras (Figure 7). (I) Heatmap of the top 30 differentially expressed genes in each of the purified graft-derived IM populations in thorax-shielded chimeras. (n=4) (J) Metascape analysis of differentially expressed genes in GMP-IM and MDP-IM isolated from thorax-shielded chimeras. (n=4) (K) Representative microscopy image of a lung lobe of a thorax-shielded chimera stained for TubB3, and statistical analysis of the distances (touching <10 microns, close <[10,100] microns, far >100 microns) of GMP- and MDP-derived cells to the closest neuron. (n=4 FOV from 2 lobes of 1 chimera)

**Supplementary Figure 7.**
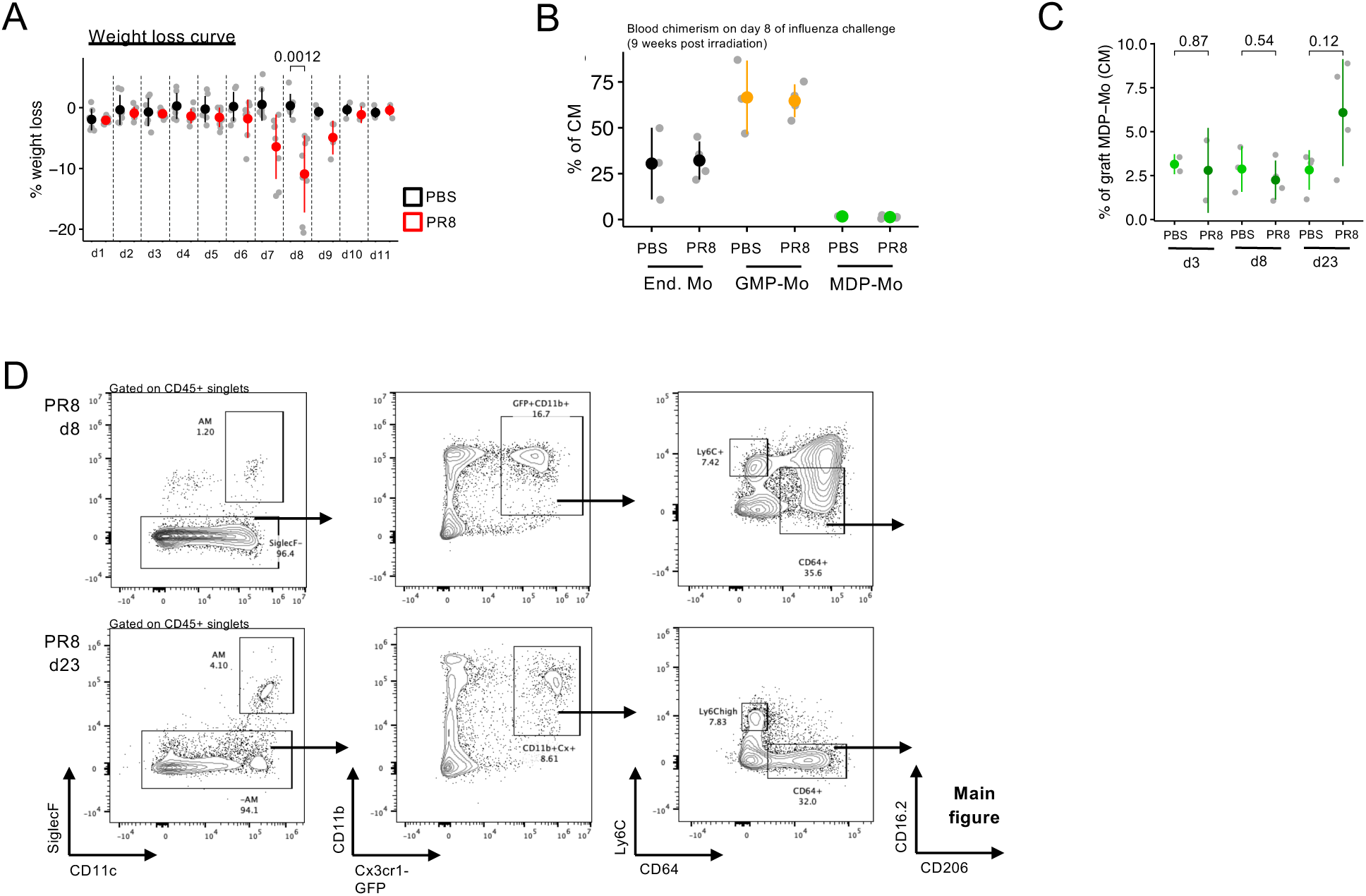
(A) Weight loss curve of PR8-infected and PBS-treated (control) thorax-shielded chimeras. (n=7-8 per condition) (B) Distribution of endogenous CM and graft-derived GMP-Mo and MDP-Mo in thorax-shielded chimeras infected with PR8 influenza or control (PBS) on d8 following challenge. (n=3-4) (C) Distribution of MDP-Mo in thorax-shielded chimeras infected with PR8 influenza or control (PBS) on several timepoints following challenge. (n=3-4 per timepoint) (D) Representative gating strategies for identification of graft-derived (GFP+) MΦ (Ly6C^−^ CD64^+^) in the lungs of thorax-shielded chimeras infected with PR8 on various timepoints following challenge. (n=4 per timepoint)

## Methods

### Mice

This study involved the following animals, all on C57BL/6 background: wild type C57BL/6 mice (Harlan); *Cx3cr1^GFP^* mice (B6.129P2(Cg)-*Cx3cr1^tm1Litt^*/J) Jax stock #005582 ^46^; *Cx3cr1^Cre^* mice (B6J.B6N(Cg)-*Cx3cr1^tm1.1(cre)/Jung^*/J) Jax stock #025524 ^11^; *Cx3cr1^DTR^* mice (B6N.129P2-*Cx3cr1^tm3(DTR)Litt^*/J) Jax stock #025629 ^48^; *Ms4a3^Cre^* mice (C57BL/6J-*Ms4a3^em2(cre)Fgnx^*/J) Jax stock #036382 ^13^; *Rosa26^LSL-TdTomato^*(B6.Cg-*Gt(ROSA)26Sor^tm9(CAG-tdTomato)Hze^*/J) Jax stock #007909 ^74^. All transgenic mice were heterozygotes for the modified alleles. Male and female mice of 7-10 weeks were used in most experiments. For generation of BM chimeras, wild type C57BL/6 mice were used as recipients. Recipient mice were lethally irradiated with a single dose of 950 cGy using an XRAD 320 machine (Precision X-Ray) and reconstituted the next day by i.v. injection of 5 x 10^6^ donor BM cells per mouse in case of whole-body irradiation. For the generation of shielded chimeras, the thoracic cavity of anesthetized mice was covered with a 6 mm-thick lead sheath. Mice were irradiated the same way, but reconstituted 6h following irradiation by i.v. injection of 5-10 x 10^6^ BM cells per mouse. Recipients were allowed to recover vor 8-10 weeks before performing experiments. All animals were maintained in a specific-pathogen-free facility with chow and water provided *ad libitum* and handled according to protocols approved by the Weizmann Institute Animal Care Committee as per international guidelines.

### LPS, CpG, IFN, Antibody, and DTx Treatments

For the LPS challenge, mice were injected intraperitoneally (i.p.) or intravenously (i.v.) with a single dose of LPS (*E. coli* O111:B4; Sigma Cat#L2630) 2.5 mg/kg. For the CpG challenge, mice were injected i.v. with a single dose of CpG (ODN 1826, Class B CpG oligonucleotide, InvivoGen Cat#tlr-1826) 5 ug/mouse and DOTAP (DOTAP liposomal transfection reagent, Sigma-Aldrich Cat#890895P-25MG) 25 ug/mouse. For the IFNg challenge, mice were injected i.v. with a single dose or i.p. for 3 consecutive days with recombinant murine IFNg (Peprotech Cat#315-05) 5 ug/mouse. To assess the CD62L involvement in NeuMo trafficking to the dura mater, recipient mice were injected 1h prior to adoptive cell transfer i.v. with purified anti-mouse CD62L antibody (clone MEL-14, BioLegend Cat#104402) 100 ug/mouse or purified rat IgG2a,k isotype ctrl antibody (clone RTK2758, BioLegend Cat#400502) 100 ug/mouse. Injection was repeated once more 24h after the initial injection. For depletion of Cx3cr1-expressing cells in *Cx3cr1^DTR^* chimeras, two initial doses of diphtheria toxin (*C. diphtheria*, Sigma Cat#D0564-1MG) 18 ng/g bodyweight per day were given before adoptive cell transfer. After cell transfer, 12 ng/g bodyweight DTx was administered every other day.

### Influenza infection

Mice were infected with a murine influenza virus (PR8 strain) diluted in PBS (phosphate buffered saline) and administered in 30μl intranasally (15μl per nostril, titer 7e4 PFU/ml / mouse). Intranasal PBS administration (15μl per nostril) served as control.

### Cell Isolation from Tissue for Flow Cytometry

Peripheral blood was collected through cardiac puncture or cheek punch. RBC were lyzed by home-made ACK buffer. Cell suspensions were then resuspended in home-made FACS buffer (2% FCS, 1 mM EDTA). Cells were stained with biotinylated lineage markers TCRb (clone H57-597), CD19 (clone 6D5), NK1.1 (clone PK136). The backbone surface panel consisted of CD11b (clone M1/70), CD115 (clone AFS98), Ly6C (clone HK1.4), CD11c (clone N418), CD319 (clone 4G2), CD177 (clone Y127), I-A/I-E (clone M5/114.15.2), CX3CR1 (clone SA011F11). Additional markers were integrated into this panel as necessary. Single cells suspensions were filtered and acquired on a Cytek Aurora 4L spectral cytometer (16V-14B-10YG-8R, Cytek Biosciences).

Lungs were perfused, tissue was collected and weighed. Tissue was transferred to digestion solution (RPMI + 1 mg/ml Col IV + 0.02 mg/ml DNase I) and minced into fine pieces. This was incubated on a shaker for 30 min at 37°C at 200 rpm. The pellet was disrupted after 15 min of incubation. After 30 min, the suspension was pipetted until homogenized and filtered through a 40 um cell strainer. The cell suspension was washed with PBS + 1mM EDTA before spinning down. The resulting pellet was then stained. Surface markers used for lung experiments included: CD45 (clone 30-F11), CD11b (clone M1/70), CD64 (clone X54-5/7.1), I-A/I-E (clone M5/114.15.2), SiglecF (clone S17007L), CD11c (clone N418), CX3CR1 (clone SA011F11), Lyve1 (clone ALY7), CD16.2 (clone 9E9), CD206 (clone C068C2). For analysis of colon and ileum, cells were isolated as previously described ^51^.

### Adoptive Cell Transfers

Femurs, tibias, and spine of donor mice were removed. BM was extracted by crushing the bones. All subsequent steps were carried out on ice. For adoptive monocyte transfers, cells were centrifuged and incubated with biotin-labeled anti-CD115 (clone AFS98, 1:200) for 20 minutes. After a washing step, cells were incubated with Streptavidin Microbeads (30-50 ul per mouse, Miltenyi Biotec). After another wash, magnetic separation was performed according to the manufacturer’s instructions. Positively selected cells (CD115+) were stained with Ly6C (clone HK1.4), CD11b (clone M1/70), CD115 (clone AFS98), and CD177 (clone Y127) and CD319 (clone 4G2) depending on th experiment. Cells were sorted on FACS Aria III (BD Biosciences) into cold RPMI + 10% FCS. Cells were spun down and counted. 2×10^5^ cells of each cell type were injected i.v. into DTx-depleted recipients. For adoptive BM precursor transfer, total CD45.2 BM cells were separated in a Ficoll gradient (Merck) and the buffy coat was collected. Cells were stained for biotinylated lineage markers (TCRb (clone H57-597), CD19 (clone 6D5), NK1.1 (clone PK136), TER-119 (clone TER-119), CD11b (clone M1/70), Ly6G (clone 1A8)), washed, and stained with secondary fluorophore-conjugated streptavidin, CD117 (clone 2B8), CD34 (clone SA376A4), Flt3 (clone A2F10), CD115 (clone AFS98), and Ly6C (clone HK1.4) antibodies. Cells were sorted on FACS Aria III (BD Biosciences) into cold RPMI + 10% FCS. Cells were spun down and counted. 20-50k of precursor cells were transferred i.v. into CD45.1 recipients.

### ETosis Assay

To induce ET formation, cells were isolated as previously described in ‘Adoptive Cell Transfers’. Freshly sorted 1 x 10^5^ cells were resuspended in RPMI medium 1640 containing 100 nM PMA, and plated on Cell-tak coated coverslips for 6 hours at 37°C. Then, medium was aspirated and cells were fixed at RT with 4% PFA for 10 min, blocked and permeabilized using PBS with 0.1% Triton X-100, 1% fetal calf serum, and 5% BSA for 30 min. Cells were then stained for citrullinated H3, MPO, and DNA (DAPI). Cover slips were imaged using an Olympus BX51 confocal laser scanning microscope, and analyzed using Fiji. 15-20 FOV for each cell subset were selected and triple-positive events were counted and normalized to all DAPI^+^ events.

### Transwell Migration Assay

Cells were isolated as detailed in ‘Adoptive Cell Transfers’ without staining and sorting steps. Importantly, for this experiment, all steps were carried out at room temperature. Transwells (24-well plate, 6.5mm diameter transwell, 5 um pore size, Costar) were pre-coated with 1% BSA at 37°C for 1 hour and then washed twice with PBS. Cells were washed twice in PBS and resuspended in binding medium (HBSS, 1% PSA, 1% HEPES, 1mM CaCl2, 2 mg/ml BSA). 100 nM of fMLP or equivalent volume of DMSO were gently resuspended in pre-warmed binding medium and 600 ul were put in the bottom well. 100 ul of cell suspension (5e6 cells/ml) were seeded in the transwell filter. Cells were let migrate for one hour at 37°C. Then, transwells were removed, medium was collected from the lower well. Cells were spun down and stained with CD11b, CD115, Ly6C, CD177, and CD319. Finally, cell suspensions were acquired on a Cytek Aurora 4L spectral cytometer.

### Phagocytosis Assay

Cells were isolated as detailed in ‘Adoptive Cell Transfers’ and incubated with GFP-expressing *Salmonella Typhimurium* in a 10:1 ratio for 2 hours at 37°C. Then, gentamycin was added for 5 minutes before spinning down. Finally, cells were acquired on a Cytek Aurora 4L spectral cytometer.

### Histology

Recipient mice were anesthetized with Pental and perfused with cold PBS. The dura mater was detached from the skull. Brain tissues were excised and fixed for 2 hours in 4% PFA at RT. Whole-mount pre-fixed tissues were blocked in 2% horse serum for 2h at RT and incubated with primary antibody overnight at 4°C. After incubation, tissues were washed thrice in PBS and incubated with secondary antibody for 2h at RT. Subsequently, tissues were incubated for 5 min with DAPI (1:10k) (Sigma), and washed thrice with PBS. Tissues were recorded with a Zeiss LSM 880 confocal laser scanning microscope with ZEN microscopy software. Image analysis was processed by Imaris software (Oxford Instruments). The following primary antibodies were used: rat monoclonal anti-CD31 (1:250, MEC13.3, Biolegend, #102502), goat polyclonal anti-CD206 (1:250, R&D Systems, #AF2535), rat monoclonal anti-MHC class II (I-A/I-E) (1:250, M5.114.15.2, Invitrogen #14-5321-82), and following secondary antibodies were used: donkey anti-rat IgG (H+L) cy3 (cat. no. 712-165-154), and donkey anti-goat IgG (H+L) Alexa Fluor 647 (cat. no. 705-605-147), which are all from Jackson Immuno Research Laboratory (JIR).

### Cubic Clearing

For cubic clearing, lungs were perfused with cold PBS. A slightly modified protocol of Matsumoto *et al* ^75^ was followed. Briefly, samples were incubated in CUBIC 1 solution : PBS (50:50) overnight at RT on a shaker. Then, samples were incubated in 100% CUBIC 1 solution for at least 3 days. Samples were then moved to CUBIC 2 solution: PBS (50:50) for overnight incubation at RT on a shaker. Following, samples were incubated in 100% CUBIC 2 solution for 2 days at 37°C on a shaker (100 rpm). After 2 days, samples were stored in mineral oil (Sigma M8410) and imaged the next day.

### CITE-seq Library Generation and Analysis

The femur was harvested from a 10 week-old C57BL6 female mouse. The BM was placed directly into ice-cold Roswell park Memorial Institute (RPMI) 1640 medium (Thermofisher). No actinomycin D was used as there were no tissue digestion steps present and all steps were performed with ice-cold buffers to maintain the cells at a low temperature. The BM was flushed by cutting the femur in two and flushing the bone cavity with a syringe with a 26g needle filled with 5% magnetic-activated cell sorting (MACS) buffer (5% inactivated fetal calf serum, 2mM EDTA (Duchefa), 1X Hanks’ buffered salt solution (HBSS) (Gibco)). This BM was flushed through a 40µm cell strainer (Corning) and the filter was washed with 5% MACS buffer. The BM was centrifuged at 450g for 6 minutes at 4°C. The supernatant was discarded, and the pellet was resuspended in home-made RBC lysis buffer. The lysis buffer was neutralized with 5% MACS buffer and the cell suspension was centrifuged. The pellet was resuspended, counted with trypan blue, and 925,000 cells were aliquoted and centrifuged. The supernatant was discarded and the cell pellet was resuspended in 25µl 1X PBS with 0.04% bovine serum albumin (BSA) buffer, containing mouse TruStain FcX (BioLegend) and the mouse cell surface protein antibody panel containing 174 oligo-conjugated antibodies (**Supplementary Table 1**). The cells were stained for 30 minutes on ice, followed by a washing step before loading. Single-cell suspensions were loaded on a GemCode Single Cell Instrument (10× Genomics) to generate single-cell gel beads in emulsion (GEM) using a GemCode Single Cell 3′ Gel Bead and Library kit v.3.1 (10x Genomics, 1000121) and a Chromium i7 Multiplex Kit (10× Genomics, 120262) as previously described ^76^. Briefly, GEM reverse-transcription incubation was performed in a 96-deep-well reaction module at 53°C for 45 min, 85°C for 5 min and ending at 4°C. Next, GEMs were broken and complementary DNA (cDNA) was cleaned up with DynaBeads MyOne Silane Beads (10x Genomics, No. 2000048) and SPRIselect Reagent Kit (Beckman Coulter, No. B23318). Full-length, barcoded cDNA was PCR amplified with a 96-deep-well reaction module at 98°C for 3 min, 11 cycles at 98°C for 15 s, 63°C for 20 s and 72°C for 1 min, followed by one cycle at 72°C for 1 min and ending at 4°C. Following cleaning up with the SPRIselect Reagent Kit and enzymatic fragmentation, library construction to generate Illumina-ready sequencing libraries was performed by the addition of R1 (read 1 primer), P5, P7, i7 sample index and R2 (read 2 primer sequence) via end-repair, A-tailing, adapter ligation, post-ligation SPRIselect cleanup/size selection and sample index PCR. The cDNA content of pre-fragmentation and post-sample index PCR samples was analyzed using the 2100 BioAnalyzer (Agilent). Sequencing libraries were loaded on an Illumina Illumina NovaSeq 6000 flow cell, with sequencing settings according to the recommendations of 10× Genomics (read 1: 26 cycles; read 2: 98 cycles; index i7: eight cycles; index i5: no cycles, pooled in a 80:20 ratio for the combined 3′ gene expression and cell surface protein samples, respectively). The Cell Ranger software (10x Genomics) was used to perform sample demultiplexing, RNA read mapping to the reference genome (mouse mm10) and RNA and ADT barcode processing, unique molecular identifiers filtering and single-cell UMI counting. The mean of the mapped RNA reads per cell was 18 864, with a sequencing saturation of 41.3%, as calculated by Cell Ranger. The ADT libraries yielded 1186 mean reads per cell, and 52.1% ADT sequencing saturation. The RNA expression matrix was further filtered and preprocessed using the Seurat (v.4.0.5) and Scater (v.1.22.0) R packages. For filtering the low-quality cell barcodes, associated with droplets that do not contain intact cells, the “emptyDrops” function of the DropletUtils package (v.1.14.2) has been applied on the RNA expression data, using an FDR cutoff of 0.01. Outlier cells were additionally identified based on three metrics (library size, number of expressed genes and mitochondrial proportion per cell); cells were tagged as outliers, when they were more than three median absolute deviations distant from the median value of each metric across all cells. Doublet score was assigned to each cell barcode based on generation of cluster-based artificial doublets with the scDblFinder function of the scDblFinder package (1.8.0). Low-abundance genes were removed using the Scater ‘calcAverage’ function and a cutoff of 0.003 mean reads per gene. The resulting RNA matrix was normalized using the global-scaling normalization and log-transformation ‘LogNormalize’ Seurat function. To mitigate the effects of cell cycle heterogeneity on the dataset, we calculating cell cycle phase scores based on canonical S and G2/M markers using the CellCycleScoring function of Seurat. Next, the cell cycle scores were regressed out from the data. Highly variable genes were detected in Seurat and the gene expression was scaled by linear transformation. Subsequently, the identified highly variable genes were used for performing principal component analysis (PCA). The PCA embeddings were used downstream for unsupervised Leiden clustering of the cells and Uniform Manifold Approximation and Projection (UMAP) dimensionality reduction as implemented in Seurat. Differential gene expression analysis was done using Wilcoxon Rank Sum test and Bonferroni correction has been applied for adjustment of the P values. Several of the identified clusters were further individually subsetted and re-clustered in order to obtain a more refined mapping of the murine BM subpopulations. During this process, we iteratively excluded cell subclusters that showed both high doublet score and expression of markers, specific for two different cell populations, e.g., of the neutrophil markers Ly6g/S100a8 and the B cell markers Cd19/Cd79a. The total number of identified cells after removing artefacts and cell doublets was 16066.The processing of the ADT expression matrix was done as described previously ^76^. In brief, the ADT cell barcodes, associated with artefact cells based on the RNA expression analysis were discarded, and the remaining data was normalized using the ASINH_GEOM transformation (inverse hyperbolic sine transformation with a cofactor).

### RNAseq Library Preparation

Peripheral blood or lung cells were isolated as previously described and sorted into 30 ul of lysis/binding buffer (home-made or Life Technologies) and stored at −80°C. mRNA was then captured with Dynabeads oligo(dT) (Life Technologies) according to the manufacturer’s guidelines. A bulk variation of MARSseq ^77^ was used for preparation of RNAseq libraries. Briefly, RNA was first reverse-transcribed with MARSseq barcoded RT primers with the Affinity Script kit (Agilent). Reverse transcription was analyzed by qRT-PCR and samples with a similar CT were pooled. Each pool was treated with Exonuclease I (NEB) for 30’ at 37°C. Double-stranded DNA was generated with the NEB second strand synthesis kit at 16°C for 2h.), Subsequently, *in vitro* transcription was performed over-night with the T7 High Yield RNA polymerase IVT kit (NEB) for 37°C overnight. Following IVT, the DNA template was removed with Turbo DNase I (Ambion) at 37°C for 15’. Amplified RNA was fragmented by incubation at 70°C for 3’ in Zn2+ RNA fragmentation reagent (Ambion). RNA was then ligated to the MARSseq ligation adaptor with T4 RNA Ligase I (NEB) at 22°C for 2h. Ligated product was reverse-transcribed using the Affinity Script RT enzyme (Agilent) and a primer complementary to the ligated product. The library was completed and amplified through a nested PCR reaction with P5_Rd1 and P7_Rd2 primers and PCR ready mix (Kappa Biosystems). Library concentrations was measured with a Qubit fluorometer (Life Technologies) and mean molecule size was determined with a 2200 TapeStation instrument. RNAseq libraries were sequenced using the Illumina NextSeq 500.

### RNAseq analysis

Data analysis was performed using the UTAP transcriptome analysis pipeline ^78^.Raw reads were trimmed using cutadapt with the parameters: -a AGATCGGAAGAGCACACGTCTGAACTCCAGTCAC -a ‘‘A–times 2 -u 3 -u 3 -q 20 -m 25). Reads were mapped to the genome (mm10, Gencode annotation version 10.0) using STAR (v2.4.2a) with the parameters –alignEndsType EndToEnd, –outFilterMismatchNoverLmax 0.05, –twopassMode Basic, –alignSoftClipAtReferenceEnds No. The pipeline quantifies the 30 of Gencode annotated genes (The 30 region contains 1,000 bases upstream of the 30 end and 100 bases downstream). UMI counting was done after marking duplicates (in-house script) using HTSeq-count in union mode. Only reads with unique mapping were considered for further analysis, and genes having minimum 5 reads in at least one sample were considered. Gene expression levels were calculated and normalized using DESeq2 with the parameters: betaPrior=True, cooksCutoff=FALSE, independentFiltering=FALSE. Batch correction was done using the *sva* (3.26.0) R when batch adjustments were required. Raw p-values were adjusted for multiple testing, using the procedure of Benjamini and Hochberg. Differentially expressed genes were selected with absolute fold change (log2)>1, and adjusted p-value <0.05. Visualization of gene expression heatmaps was done using Gene-e, using log normalized values. Clustering was applied on log2 transformed and standardized expression values, using the k-means algorithm (Euclidian method). For Gene Ontology (GO) term analysis, Metascape was used ^57^, and all heatmaps were made by Morpheus, https://software.broadinstitute.org/morpheus.

### Image analysis

Images of lung sections of thorax-shielded [ *Ms4a3^Cre^:R26^LSL-TdTom^:Cx3cr1^gfp^* > WT] chimeras stined with anti-Tubb3 antibody were analyzed with open-source software using Fiji ^79^, Ilastik ^80^ and Cellpose ^81^. Specifically, we used out-of-the-box Cellpose’s “cyto2” model to identify cells: We used cell diameter = 20pixels for identifying the GMP-derived MΦ, and cell diameter = 10pixels for identifying the MDP-derived MΦ. To segment the neurons, we trained an Ilastik model using all 4 images on which the distance analysis was performed. We used Fiji’s distance transform plugin to extract the Mean and Min. distance between the segmented neurons and the identified cells and MΦ. The Fiji macro used to run the analysis, together with the trained Ilastik model are deposited and available for download on GitHub (link to be provided upon paper acceptance).

### Statistical Analysis

In all experiment, data are presented as mean +− standard deviation (SD). Statistical tests were selected based on appropriate assumptions with respect to data distribution and variance characteristics. Student’s t test (two-tailed) was applied to demonstrate statistical differences between two groups, and Two-way ANOVA was used when appropriate. Sample sizes were chosen according to standard guidelines. Number of animals is indicated as ‘‘n’’ and presented with the number of dots on the graphs.

## References

1. Guilliams, M., Mildner, A., and Yona, S. (2018). Developmental and Functional Heterogeneity of Monocytes. Immunity 49, 595–613. 10.1016/j.immuni.2018.10.005.

2. Hettinger, J., Richards, D.M., Hansson, J., Barra, M.M., Joschko, A.-C., Krijgsveld, J., and Feuerer, M. (2013). Origin of monocytes and macrophages in a committed progenitor. Nat Immunol 14, ni.2638. 10.1038/ni.2638.

3. Yáñez, A., Ng, M.Y., Hassanzadeh-Kiabi, N., and Goodridge, H.S. (2015). IRF8 acts in lineage-committed rather than oligopotent progenitors to control neutrophil vs monocyte production. Blood 125, 1452–1459. 10.1182/blood-2014-09-600833.

4. Ginhoux, F., and Jung, S. (2014). Monocytes and macrophages: developmental pathways and tissue homeostasis. Nat Rev Immunol 14, 392–404. 10.1038/nri3671.

5. Passlick, B., Flieger, D., and Ziegler-Heitbrock, H. (1989). Identification and characterization of a novel monocyte subpopulation in human peripheral blood. Blood 74, 2527–2534. 10.1182/blood.v74.7.2527.bloodjournal7472527.

6. Palframan, R.T., Jung, S., Cheng, G., Weninger, W., Luo, Y., Dorf, M., Littman, D.R., Rollins, B.J., Zweerink, H., Rot, A., et al. (2001). Inflammatory chemokine transport and presentation in HEV: a remote control mechanism for monocyte recruitment to lymph nodes in inflamed tissues. The Journal of experimental medicine 194, 1361–1373.

7. Geissmann, F., Jung, S., and Littman, D.R. (2003). Blood monocytes consist of two principal subsets with distinct migratory properties. Immunity 19, 71–82. 10.1016/s1074-7613(03)00174-2.

8. Trzebanski, S., and Jung, S. (2020). Plasticity of Monocyte Development and Monocyte Fates. Immunol Lett 227, 66–78. 10.1016/j.imlet.2020.07.007.

9. Auffray, C., Fogg, D., Garfa, M., Elain, G., Join-Lambert, O., Kayal, S., Sarnacki, S., Cumano, A., Lauvau, G., and Geissmann, F. (2007). Monitoring of Blood Vessels and Tissues by a Population of Monocytes with Patrolling Behavior. Science 317, 666–670. 10.1126/science.1142883.

10. Mildner, A., Schönheit, J., Giladi, A., David, E., Lara-Astiaso, D., Lorenzo-Vivas, E., Paul, F., Chappell-Maor, L., Priller, J., Leutz, A., et al. (2017). Genomic Characterization of Murine Monocytes Reveals C/EBPβ Transcription Factor Dependence of Ly6C− Cells. Immunity 46, 849–862.e7. 10.1016/j.immuni.2017.04.018.

11. Yona, S., Kim, K.-W., Wolf, Y., Mildner, A., Varol, D., Breker, M., Strauss-Ayali, D., Viukov, S., Guilliams, M., Misharin, A., et al. (2013). Fate Mapping Reveals Origins and Dynamics of Monocytes and Tissue Macrophages under Homeostasis. Immunity 38, 79–91. 10.1016/j.immuni.2012.12.001.

12. Patel, A.A., Zhang, Y., Fullerton, J.N., Boelen, L., Rongvaux, A., Maini, A.A., Bigley, V., Flavell, R.A., Gilroy, D.W., Asquith, B., et al. (2017). The fate and lifespan of human monocyte subsets in steady state and systemic inflammation. The Journal of experimental medicine 74, jem.20170355–11. 10.1084/jem.20170355.

13. Liu, Z., Gu, Y., Chakarov, S., Bleriot, C., Kwok, I., Chen, X., Shin, A., Huang, W., Dress, R.J., Dutertre, C.-A., et al. (2019). Fate Mapping via Ms4a3-Expression History Traces Monocyte-Derived Cells. Cell 178, 1509–1525.e19. 10.1016/j.cell.2019.08.009.

14. Menezes, S., Melandri, D., Anselmi, G., Perchet, T., Loschko, J., Dubrot, J., Patel, R., Gautier, E.L., Hugues, S., Longhi, M.P., et al. (2016). The Heterogeneity of Ly6Chi Monocytes Controls Their Differentiation into iNOS+ Macrophages or Monocyte-Derived Dendritic Cells. Immunity 45, 1205–1218. 10.1016/j.immuni.2016.12.001.

15. Briseño, C.G., Haldar, M., Kretzer, N.M., Wu, X., Theisen, D.J., KC, W., Durai, V., Grajales-Reyes, G.E., Iwata, A., Bagadia, P., et al. (2016). Distinct Transcriptional Programs Control Cross-Priming in Classical and Monocyte-Derived Dendritic Cells. Cell Rep. 15, 2462–2474. 10.1016/j.celrep.2016.05.025.

16. Cheong, C., Matos, I., Choi, J.-H., Dandamudi, D.B., Shrestha, E., Longhi, M.P., Jeffrey, K.L., Anthony, R.M., Kluger, C., Nchinda, G., et al. (2010). Microbial Stimulation Fully Differentiates Monocytes to DC-SIGN/CD209+ Dendritic Cells for Immune T Cell Areas. Cell 143, 416–429. 10.1016/j.cell.2010.09.039.

17. Auffray, C., Fogg, D.K., Narni-Mancinelli, E., Senechal, B., Trouillet, C., Saederup, N., Leemput, J., Bigot, K., Campisi, L., Abitbol, M., et al. (2009). CX3CR1+ CD115+ CD135+ common macrophage/DC precursors and the role of CX3CR1 in their response to inflammation. J. Exp. Med. 206, 595–606. 10.1084/jem.20081385.

18. Varol, C., Landsman, L., Fogg, D.K., Greenshtein, L., Gildor, B., Margalit, R., Kalchenko, V., Geissmann, F., and Jung, S. (2007). Monocytes give rise to mucosal, but not splenic, conventional dendritic cells. J Exp Medicine 204, 171–180. 10.1084/jem.20061011.

19. Gamrekelashvili, J., Giagnorio, R., Jussofie, J., Soehnlein, O., Duchene, J., Briseño, C.G., Ramasamy, S.K., Krishnasamy, K., Limbourg, A., Häger, C., et al. (2016). Regulation of monocyte cell fate by blood vessels mediated by Notch signalling. Nat. Commun. 7, 12597. 10.1038/ncomms12597.

20. Gamrekelashvili, J., Kapanadze, T., Sablotny, S., Ratiu, C., Dastagir, K., Lochner, M., Karbach, S., Wenzel, P., Sitnow, A., Fleig, S., et al. (2020). Notch and TLR signaling coordinate monocyte cell fate and inflammation. eLife 9, e57007. 10.7554/elife.57007.

21. Carlin, L.M., Stamatiades, E.G., Auffray, C., Hanna, R.N., Glover, L., Vizcay-Barrena, G., Hedrick, C.C., Cook, H.T., Diebold, S., and Geissmann, F. (2013). Nr4a1-Dependent Ly6Clow Monocytes Monitor Endothelial Cells and Orchestrate Their Disposal. Cell 153, 362–375. 10.1016/j.cell.2013.03.010.

22. Landsman, L., Bar-On, L., Zernecke, A., Kim, K.-W., Krauthgamer, R., Shagdarsuren, E., Lira, S.A., Weissman, I.L., Weber, C., and Jung, S. (2009). CX3CR1 is required for monocyte homeostasis and atherogenesis by promoting cell survival. Blood 113, 963–972. 10.1182/blood-2008-07-170787.

23. Schyns, J., Bai, Q., Ruscitti, C., Radermecker, C., Schepper, S.D., Chakarov, S., Farnir, F., Pirottin, D., Ginhoux, F., Boeckxstaens, G., et al. (2019). Non-classical tissue monocytes and two functionally distinct populations of interstitial macrophages populate the mouse lung. Nat Commun 10, 3964. 10.1038/s41467-019-11843-0.

24. Evren, E., Ringqvist, E., Tripathi, K.P., Sleiers, N., Rives, I.C., Alisjahbana, A., Gao, Y., Sarhan, D., Halle, T., Sorini, C., et al. (2020). Distinct developmental pathways from blood monocytes generate human lung macrophage diversity. Immunity. 10.1016/j.immuni.2020.12.003.

25. Hoffman, D., Tevet, Y., Trzebanski, S., Rosenberg, G., Vainman, L., Solomon, A., Hen-Avivi, S., Ben-Moshe, N.B., and Avraham, R. (2021). A non-classical monocyte-derived macrophage subset provides a splenic replication niche for intracellular Salmonella. Immunity 54, 2712–2723.e6. 10.1016/j.immuni.2021.10.015.

26. Yáñez, A., Coetzee, S.G., Olsson, A., Muench, D.E., Berman, B.P., Hazelett, D.J., Salomonis, N., Grimes, H.L., and Goodridge, H.S. (2017). Granulocyte-Monocyte Progenitors and Monocyte-Dendritic Cell Progenitors Independently Produce Functionally Distinct Monocytes. Immunity 47, 890–902.e4. 10.1016/j.immuni.2017.10.021.

27. Weinreb, C., Rodriguez-Fraticelli, A., Camargo, F.D., and Klein, A.M. (2020). Lineage tracing on transcriptional landscapes links state to fate during differentiation. Science, eaaw3381. 10.1126/science.aaw3381.

28. Fogg, D.K., Sibon, C., Miled, C., Jung, S., Aucouturier, P., Littman, D.R., Cumano, A., and Geissmann, F. (2006). A Clonogenic Bone Marrow Progenitor Specific for Macrophages and Dendritic Cells. Science 311, 83–87. 10.1126/science.1117729.

29. Ikeda, N., Kubota, H., Suzuki, R., Morita, M., Yoshimura, A., Osada, Y., Kishida, K., Kitamura, D., Iwata, A., Yotsumoto, S., et al. (2023). The early neutrophil-committed progenitors aberrantly differentiate into immunoregulatory monocytes during emergency myelopoiesis. Cell Reports 42, 112165. 10.1016/j.celrep.2023.112165.

30. Anderson, D.A., Dutertre, C.-A., Ginhoux, F., and Murphy, K.M. (2020). Genetic models of human and mouse dendritic cell development and function. Nat Rev Immunol, 1–15. 10.1038/s41577-020-00413-x.

31. Meredith, M.M., Liu, K., Kamphorst, A.O., Idoyaga, J., Yamane, A., Guermonprez, P., Rihn, S., Yao, K.-H., Silva, I.T., Oliveira, T.Y., et al. (2012). Zinc finger transcription factor zDC is a negative regulator required to prevent activation of classical dendritic cells in the steady state. J Exp Med 209, 1583–1593. 10.1084/jem.20121003.

32. Satpathy, A.T., KC, W., Albring, J.C., Edelson, B.T., Kretzer, N.M., Bhattacharya, D., Murphy, T.L., and Murphy, K.M. (2012). Zbtb46 expression distinguishes classical dendritic cells and their committed progenitors from other immune lineages. J Exp Med 209, 1135–1152. 10.1084/jem.20120030.

33. Chong, S.Z., Evrard, M., Devi, S., Chen, J., Lim, J.Y., See, P., Zhang, Y., Adrover, J.M., Lee, B., Tan, L., et al. (2016). CXCR4 identifies transitional bone marrow premonocytes that replenish the mature monocyte pool for peripheral responses. J Exp Med 213, 2293–2314. 10.1084/jem.20160800.

34. Steppich, B., Dayyani, F., Gruber, R., Lorenz, R., Mack, M., and Ziegler-Heitbrock, H.W.L. (2000). Selective mobilization of CD14+CD16+ monocytes by exercise. Am. J. Physiol.-Cell Physiol. 279, C578–C586. 10.1152/ajpcell.2000.279.3.c578.

35. Janssen, H., Kahles, F., Liu, D., Downey, J., Koekkoek, L.L., Roudko, V., D’Souza, D., McAlpine, C.S., Halle, L., Poller, W.C., et al. (2023). Monocytes re-enter the bone marrow during fasting and alter the host response to infection. Immunity 56, 783–796.e7. 10.1016/j.immuni.2023.01.024.

36. Askenase, M.H., Han, S.-J., Byrd, A.L., Morais da Fonseca, D., Bouladoux, N., Wilhelm, C., Konkel, J.E., Hand, T.W., Lacerda-Queiroz, N., Su, X., et al. (2015). Bone-Marrow-Resident NK Cells Prime Monocytes for Regulatory Function during Infection. Immunity 42, 1130–1142. 10.1016/j.immuni.2015.05.011.

37. Biram, A., Liu, J., Hezroni, H., Davidzohn, N., Schmiedel, D., Khatib-Massalha, E., Haddad, M., Grenov, A., Lebon, S., Salame, T.M., et al. (2022). Bacterial infection disrupts established germinal center reactions through monocyte recruitment and impaired metabolic adaptation. Immunity 55, 442–458.e8. 10.1016/j.immuni.2022.01.013.

38. Papayannopoulos, V. (2018). Neutrophil extracellular traps in immunity and disease. Nat Rev Immunol 18, 134–147. 10.1038/nri.2017.105.

39. Kolaczkowska, E., and Kubes, P. (2013). Neutrophil recruitment and function in health and inflammation. Nat. Rev. Immunol. 13, 159–175. 10.1038/nri3399.

40. Castanheira, F.V.S., and Kubes, P. (2019). Neutrophils and NETs in modulating acute and chronic inflammation. Blood 133, 2178–2185. 10.1182/blood-2018-11-844530.

41. Hartt, J.K., Barish, G., Murphy, P.M., and Gao, J.-L. (1999). N-Formylpeptides Induce Two Distinct Concentration Optima for Mouse Neutrophil Chemotaxis by Differential Interaction with Two N-Formylpeptide Receptor (Fpr) Subtypes. J Exp Medicine 190, 741–748. 10.1084/jem.190.5.741.

42. Li, P., Li, M., Lindberg, M.R., Kennett, M.J., Xiong, N., and Wang, Y. (2010). PAD4 is essential for antibacterial innate immunity mediated by neutrophil extracellular traps. J Exp Med 207, 1853–1862. 10.1084/jem.20100239.

43. Granger, V., Faille, D., Marani, V., Noël, B., Gallais, Y., Szely, N., Flament, H., Pallardy, M., Chollet-Martin, S., and Chaisemartin, L. (2017). Human blood monocytes are able to form extracellular traps. J. Leukoc. Biol. 102, 775–781. 10.1189/jlb.3ma0916-411r.

44. Ginhoux, F., and Guilliams, M. (2016). Tissue-Resident Macrophage Ontogeny and Homeostasis. Immunity 44, 439–449. 10.1016/j.immuni.2016.02.024.

45. Mildner, A., Yona, S., and Jung, S. (2013). Chapter Three A Close Encounter of the Third Kind Monocyte-Derived Cells. Adv. Immunol. 120, 69–103. 10.1016/b978-0-12-417028-5.00003-x.

46. Jung, S., Aliberti, J., Graemmel, P., Sunshine, M.J., Kreutzberg, G.W., Sher, A., and Littman, D.R. (2000). Analysis of Fractalkine Receptor CX3CR1 Function by Targeted Deletion and Green Fluorescent Protein Reporter Gene Insertion. Mol. Cell. Biol. 20, 4106–4114. 10.1128/mcb.20.11.4106-4114.2000.

47. Varol, C., Vallon-Eberhard, A., Elinav, E., Aychek, T., Shapira, Y., Luche, H., Fehling, H.J., Hardt, W.-D., Shakhar, G., and Jung, S. (2009). Intestinal Lamina Propria Dendritic Cell Subsets Have Different Origin and Functions. Immunity 31, 502–512. 10.1016/j.immuni.2009.06.025.

48. Diehl, G.E., Longman, R.S., Zhang, J.-X., Breart, B., Galan, C., Cuesta, A., Schwab, S.R., and Littman, D.R. (2013). Microbiota restricts trafficking of bacteria to mesenteric lymph nodes by CX3CR1hi cells. Nature 494, 116–120. 10.1038/nature11809.

49. Aychek, T., Mildner, A., Yona, S., Kim, K.-W., Lampl, N., Reich-Zeliger, S., Boon, L., Yogev, N., Waisman, A., Cua, D.J., et al. (2015). IL-23-mediated mononuclear phagocyte crosstalk protects mice from Citrobacter rodentium-induced colon immunopathology. Nat. Commun. 6, 6525. 10.1038/ncomms7525.

50. Bianchini, M., Duchêne, J., Santovito, D., Schloss, M.J., Evrard, M., Winkels, H., Aslani, M., Mohanta, S.K., Horckmans, M., Blanchet, X., et al. (2019). PD-L1 expression on nonclassical monocytes reveals their origin and immunoregulatory function. Sci Immunol 4, eaar3054. 10.1126/sciimmunol.aar3054.

51. Gross-Vered, M., Trzebanski, S., Shemer, A., Bernshtein, B., Curato, C., Stelzer, G., Salame, T.-M., David, E., Boura-Halfon, S., Chappell-Maor, L., et al. (2020). Defining murine monocyte differentiation into colonic and ileal macrophages. Elife 9, e49998. 10.7554/elife.49998.

52. Goldmann, T., Wieghofer, P., Jordão, M.J.C., Prutek, F., Hagemeyer, N., Frenzel, K., Amann, L., Staszewski, O., Kierdorf, K., Krueger, M., et al. (2016). Origin, fate and dynamics of macrophages at central nervous system interfaces. Nat Immunol 17, 797–805. 10.1038/ni.3423.

53. Hove, H.V., Martens, L., Scheyltjens, I., Vlaminck, K.D., Antunes, A.R.P., Prijck, S.D., Vandamme, N., Schepper, S.D., Isterdael, G.V., Scott, C.L., et al. (2019). A single-cell atlas of mouse brain macrophages reveals unique transcriptional identities shaped by ontogeny and tissue environment. Nat. Neurosci. 22, 1021–1035. 10.1038/s41593-019-0393-4.

54. Shemer, A., Grozovski, J., Tay, T.L., Tao, J., Volaski, A., Süß, P., Ardura-Fabregat, A., Gross-Vered, M., Kim, J.-S., David, E., et al. (2018). Engrafted parenchymal brain macrophages differ from microglia in transcriptome, chromatin landscape and response to challenge. Nat. Commun. 9, 5206. 10.1038/s41467-018-07548-5.

55. Gibbings, S.L., Thomas, S.M., Atif, S.M., McCubbrey, A.L., Desch, A.N., Danhorn, T., Leach, S.M., Bratton, D.L., Henson, P.M., Janssen, W.J., et al. (2017). Three Unique Interstitial Macrophages in the Murine Lung at Steady State. Am. J. Respir. Cell Mol. Biology 57, 66–76. 10.1165/rcmb.2016-0361oc.

56. Vanneste, D., Bai, Q., Hasan, S., Peng, W., Pirottin, D., Schyns, J., Maréchal, P., Ruscitti, C., Meunier, M., Liu, Z., et al. (2023). MafB-restricted local monocyte proliferation precedes lung interstitial macrophage differentiation. Nat. Immunol. 24, 827–840. 10.1038/s41590-023-01468-3.

57. Zhou, Y., Zhou, B., Pache, L., Chang, M., Khodabakhshi, A.H., Tanaseichuk, O., Benner, C., and Chanda, S.K. (2019). Metascape provides a biologist-oriented resource for the analysis of systems-level datasets. Nat Commun 10, 1523. 10.1038/s41467-019-09234-6.

58. Chakarov, S., Lim, H.Y., Tan, L., Lim, S.Y., See, P., Lum, J., Zhang, X.-M., Foo, S., Nakamizo, S., Duan, K., et al. (2019). Two distinct interstitial macrophage populations coexist across tissues in specific subtissular niches. Science 363, eaau0964. 10.1126/science.aau0964.

59. Zhang, R., Ghosh, S.N., Zhu, D., North, P.E., Fish, B.L., Morrow, N.V., Lowry, T., Nanchal, R., Jacobs, E.R., Moulder, J.E., et al. (2008). Structural and functional alterations in the rat lung following whole thoracic irradiation with moderate doses: Injury and recovery. Int. J. Radiat. Biol. 84, 487–497. 10.1080/09553000802078396.

60. Ghosh, S.N., Wu, Q., Mäder, M., Fish, B.L., Moulder, J.E., Jacobs, E.R., Medhora, M., and Molthen, R.C. (2009). Vascular Injury After Whole Thoracic X-Ray Irradiation in the Rat. Int. J. Radiat. Oncol.Biol.Phys. 74, 192–199. 10.1016/j.ijrobp.2009.01.006.

61. Janssen, W.J., Muldrow, A., Kearns, M.T., Barthel, L., and Henson, P.M. (2010). Development and characterization of a lung-protective method of bone marrow transplantation in the mouse. J. Immunol. Methods 357, 1–9. 10.1016/j.jim.2010.03.013.

62. van de Laar, L., Saelens, W., De Prijck, S., Martens, L., Scott, C.L., Van Isterdael, G., Hoffmann, E., Beyaert, R., Saeys, Y., Lambrecht, B.N., et al. (2016). Yolk Sac Macrophages, Fetal Liver, and Adult Monocytes Can Colonize an Empty Niche and Develop into Functional Tissue-Resident Macrophages. Immunity 44, 755–768. 10.1016/j.immuni.2016.02.017.

63. Iliakis, C.S., Kulikauskaite, J., Aegerter, H., Li, F., Piattini, F., Jakubzick, C.V., Guilliams, M., Kopf, M., and Wack, A. (2023). The role of recruitment versus training in influenza-induced lasting changes to alveolar macrophage function. Nat. Immunol. 24, 1639–1641. 10.1038/s41590-023-01602-1.

64. Bain, C.C., Bravo-Blas, A., Scott, C.L., Perdiguero, E.G., Geissmann, F., Henri, S., Malissen, B., Osborne, L.C., Artis, D., and Mowat, A.M. (2014). Constant replenishment from circulating monocytes maintains the macrophage pool in the intestine of adult mice. Nat. Immunol. 15, 929– 937. 10.1038/ni.2967.

65. Kim, J.-S., Trzebanski, S., Shin, S.-H., Ilani, N.C., Kaushansky, N., Scheller, M., Solomon, A., Liu, Z., Aust, O., Boura-Halfon, S., et al. (2023). Monocyte-derived microglia with Dnmt3a mutation cause motor pathology in aging mice. bioRxiv, 2023.11.16.567402. 10.1101/2023.11.16.567402.

66. Evans, M.A., Sano, S., and Walsh, K. (2019). Cardiovascular Disease, Aging, and Clonal Hematopoiesis. Annu. Rev. Pathol.: Mech. Dis. 15, 1–20. 10.1146/annurev-pathmechdis-012419-032544.

67. Arends, C.M., Liman, T.G., Strzelecka, P.M., Kufner, A., Löwe, P., Huo, S., Stein, C.M., Piper, S.K., Tilgner, M., Sperber, P.S., et al. (2022). Associations of clonal hematopoiesis with recurrent vascular events and death in patients with incident ischemic stroke. Blood 141, 787– 799. 10.1182/blood.2022017661.

68. Xie, Q., Klesney-Tait, J., Keck, K., Parlet, C., Borcherding, N., Kolb, R., Li, W., Tygrett, L., Waldschmidt, T., Olivier, A., et al. (2015). Characterization of a novel mouse model with genetic deletion of CD177. Protein Cell 6, 117–126. 10.1007/s13238-014-0109-1.

69. Amorim, A., Feo, D.D., Friebel, E., Ingelfinger, F., Anderfuhren, C.D., Krishnarajah, S., Andreadou, M., Welsh, C.A., Liu, Z., Ginhoux, F., et al. (2022). IFNγ and GM-CSF control complementary differentiation programs in the monocyte-to-phagocyte transition during neuroinflammation. Nat. Immunol. 23, 217–228. 10.1038/s41590-021-01117-7.

70. Giladi, A., Wagner, L.K., Li, H., Dörr, D., Medaglia, C., Paul, F., Shemer, A., Jung, S., Yona, S., Mack, M., et al. (2020). Cxcl10+ monocytes define a pathogenic subset in the central nervous system during autoimmune neuroinflammation. Nat. Immunol. 21, 525–534. 10.1038/s41590-020-0661-1.

71. Simmons, D.P., Nguyen, H.N., Gomez-Rivas, E., Jeong, Y., Jonsson, A.H., Chen, A.F., Lange, J.K., Dyer, G.S., Blazar, P., Earp, B.E., et al. (2022). SLAMF7 engagement superactivates macrophages in acute and chronic inflammation. Sci. Immunol. 7, eabf2846. 10.1126/sciimmunol.abf2846.

72. Goodridge, H.S. (2023). Aging of classical monocyte subsets. Aging (Albany NY) 15, 290–292. 10.18632/aging.204493.

73. Barman, P.K., Shin, J.E., Lewis, S.A., Kang, S., Wu, D., Wang, Y., Yang, X., Nagarkatti, P.S., Nagarkatti, M., Messaoudi, I., et al. (2022). Production of MHCII-expressing classical monocytes increases during aging in mice and humans. Aging Cell 21, e13701. 10.1111/acel.13701.

74. Madisen, L., Zwingman, T.A., Sunkin, S.M., Oh, S.W., Zariwala, H.A., Gu, H., Ng, L.L., Palmiter, R.D., Hawrylycz, M.J., Jones, A.R., et al. (2010). A robust and high-throughput Cre reporting and characterization system for the whole mouse brain. Nat. Neurosci. 13, 133–140. 10.1038/nn.2467.

75. Matsumoto, K., Mitani, T.T., Horiguchi, S.A., Kaneshiro, J., Murakami, T.C., Mano, T., Fujishima, H., Konno, A., Watanabe, T.M., Hirai, H., et al. (2019). Advanced CUBIC tissue clearing for whole-organ cell profiling. Nat. Protoc. 14, 3506–3537. 10.1038/s41596-019-0240-9.

76. Scheyltjens, I., Hove, H.V., Vlaminck, K.D., Kancheva, D., Bastos, J., Vara-Pérez, M., Antunes, A.R.P., Martens, L., Scott, C.L., Ginderachter, J.A.V., et al. (2022). Single-cell RNA and protein profiling of immune cells from the mouse brain and its border tissues. Nat. Protoc. 17, 2354–2388. 10.1038/s41596-022-00716-4.

77. Jaitin, D.A., Kenigsberg, E., Keren-Shaul, H., Elefant, N., Paul, F., Zaretsky, I., Mildner, A., Cohen, N., Jung, S., Tanay, A., et al. (2014). Massively Parallel Single-Cell RNA-Seq for Marker-Free Decomposition of Tissues into Cell Types. Science 343, 776–779. 10.1126/science.1247651.

78. Kohen, R., Barlev, J., Hornung, G., Stelzer, G., Feldmesser, E., Kogan, K., Safran, M., and Leshkowitz, D. (2019). UTAP: User-friendly Transcriptome Analysis Pipeline. BMC Bioinform. 20, 154. 10.1186/s12859-019-2728-2.

79. Schindelin, J., Arganda-Carreras, I., Frise, E., Kaynig, V., Longair, M., Pietzsch, T., Preibisch, S., Rueden, C., Saalfeld, S., Schmid, B., et al. (2012). Fiji: an open-source platform for biological-image analysis. Nat. Methods 9, 676–682. 10.1038/nmeth.2019.

80. Berg, S., Kutra, D., Kroeger, T., Straehle, C.N., Kausler, B.X., Haubold, C., Schiegg, M., Ales, J., Beier, T., Rudy, M., et al. (2019). ilastik: interactive machine learning for (bio)image analysis. Nat. Methods 16, 1226–1232. 10.1038/s41592-019-0582-9.

81. Stringer, C., Wang, T., Michaelos, M., and Pachitariu, M. (2021). Cellpose: a generalist algorithm for cellular segmentation. Nat. Methods 18, 100–106. 10.1038/s41592-020-01018-x.

