## Supplementary material for "Murine GMP- and MDP-derived classical monocytes have distinct functions and fates": Suppl Table 1

| Barcode | TotalSeq ID | Reactivity | Isotype |
|---|---|---|---|
| 4-1BB (CD13 | A0890 | Mouse | Rat IgG2a, κ |
| CD102 | A0104 | Mouse | Rat IgG2a, κ |
| CD103 | A0201 | Mouse | Armenian Hamster IgG |
| CD105 | A0812 | Mouse | Rat IgG2a, κ |
| CD106 | A0226 | Mouse | Rat IgG2a, κ |
| CD107a | A0905 | Mouse | Rat IgG2a, κ |
| CD115 | A0105 | Mouse | Rat IgG2a, κ |
| CD117 | A0012 | Mouse | Rat IgG2b, κ |
| CD11a | A0595 | Mouse | Rat IgG2a, κ |
| CD11b | A0014 | Human, Mouse | Rat IgG2b, κ |
| CD11c | A0106 | Mouse | Armenian Hamster IgG |
| CD122 | A0227 | Mouse | Rat IgG2a, κ |
| CD124 | A0916 | Mouse | Rat IgG2b, κ |
| CD127 | A0198 | Mouse | Rat IgG2a, κ |
| CD134 | A0195 | Mouse | Rat IgG1, κ |
| CD135 | A0098 | Mouse | Rat IgG2a, κ |
| CD137 | A0194 | Mouse | Syrian Hamster IgG |
| CD138 | A0810 | Mouse | Rat IgG2a, κ |
| CD14 | A0424 | Mouse | Rat IgG2a, κ |
| CD140a | A0573 | Mouse | Rat IgG2a, κ |
| CD146 | A0134 | Human | Mouse IgG1, κ |
| CD15 | A0076 | Human, Mouse | Mouse IgM, κ |
| CD150 | A0203 | Mouse | Rat IgG2a, λ |
| CD152 | A0388 | Mouse | Armenian Hamster IgG |
| CD16-CD32 | A0109 | Mouse | Rat IgG2a, λ |
| CD163 | A0417 | Mouse | Rat IgG2a, κ |
| CD169 | A0440 | Mouse | Rat IgG2a, κ |
| CD172a | A0422 | Mouse | Rat IgG1, κ |
| CD183 | A0228 | Mouse | Armenian Hamster IgG |
| CD185 | A0846 | Mouse | Rat IgG2b, κ |
| CD19 | A0093 | Mouse | Rat IgG2a, κ |
| CD192 | A0426 | Mouse | Rat IgG2b, κ |
| CD193 | A0808 | Mouse | Rat IgG2a, κ |
| CD195 | A0376 | Mouse | Armenian Hamster IgG |
| CD196 | A0225 | Mouse | Armenian Hamster IgG |
| CD197 | A0377 | Mouse | Rat IgG2a, κ |
| CD1d | A0851 | Mouse | Rat IgG2b, κ |
| CD2 | A0892 | Mouse | Rat IgG2b, λ |
| CD20 | A0192 | Mouse | Rat IgG2b, κ |
| CD200 | A0079 | Mouse | Rat IgG2a, κ |
| CD200R | A0807 | Mouse | Rat IgG2a, κ |
| CD200R3 | A0809 | Mouse | Rat IgG2a, κ |
| CD201 | A0439 | Mouse | Rat IgG2a, κ |
| CD204 | A0448 | Mouse | Rat IgG2a |
| CD206 | A0173 | Mouse | Rat IgG2a, κ |
| CD207 | A0437 | Human, Mouse | Mouse IgG2a, κ |
| CD21-CD35 | A0107 | Mouse | Rat IgG2a, κ |

| | | | |
|---|---|---|---|
| CD22 | A0827 | Mouse | Rat IgG1, κ |
| CD223 | A0378 | Mouse | Rat IgG1, κ |
| CD226 | A0852 | Mouse | Rat IgG2b, κ |
| CD23 | A0108 | Mouse | Rat IgG2a, κ |
| CD24 | A0212 | Mouse | Rat IgG2b, κ |
| CD25 | A0097 | Mouse | Rat IgG1, λ |
| CD26 | A0883 | Mouse | Rat IgG2a, κ |
| CD27 | A0191 | Human, Mouse, Rat | Armenian Hamster IgG |
| CD270 | A0885 | Mouse | Armenian Hamster IgG |
| CD272 | A0881 | Mouse | Armenian Hamster IgG |
| CD274 | A0190 | Mouse | Rat IgG2a, κ |
| CD278 | A0171 | Human, Mouse, Rat | Armenian Hamster IgG |
| CD279 | A0004 | Mouse | Rat IgG2b, κ |
| CD28 | A0204 | Mouse | Syrian Hamster IgG |
| CD29 | A0570 | Mouse, Rat | Armenian Hamster IgG |
| CD3 | A0182 | Mouse | Rat IgG2b, κ |
| CD300c_d | A0876 | Mouse | Rat IgG2b, κ |
| CD300LG | A0416 | Mouse | Rat IgG2a, κ |
| CD301a | A0551 | Mouse | Rat IgG2a, κ |
| CD301b | A0566 | Mouse | Rat IgG2a, λ |
| CD304 | A0552 | Mouse | Rat IgG2a, κ |
| CD309 | A0554 | Mouse | Rat IgG2a, κ |
| CD31 | A0904 | Mouse | Rat IgG2a, κ |
| CD314 | A0835 | Mouse | Rat IgG1, κ |
| CD317 | A0811 | Mouse | Rat IgG2b, κ |
| CD326 | A0449 | Mouse | Rat IgG2a, κ |
| CD335 | A0184 | Mouse | Rat IgG2a, κ |
| CD34 | A0857 | Mouse | Armenian Hamster IgG |
| CD357 | A0193 | Mouse | Rat IgG2b, λ |
| CD36 | A0555 | Mouse | Armenian Hamster IgG |
| CD366 | A0003 | Mouse | Rat IgG2a, κ |
| CD370 | A0556 | Mouse | Rat IgG1, κ |
| CD371 | A0825 | Mouse | Rat IgG2a, κ |
| CD38 | A0557 | Mouse | Rat IgG2a, κ |
| CD39 | A0834 | Mouse | Rat IgG2a, κ |
| CD4 | A0001 | Mouse | Rat IgG2a, κ |
| CD41 | A0443 | Mouse | Rat IgG1, κ |
| CD43 | A0110 | Mouse | Rat IgG2b |
| CD45R-B220 | A0103 | Human, Mouse | Rat IgG2a, κ |
| CD48 | A0429 | Mouse | Armenian Hamster IgG |
| CD49a | A0850 | Mouse | Armenian Hamster IgG |
| CD49b | A0421 | Mouse | Armenian Hamster IgG |
| CD49d | A0078 | Mouse | Rat IgG2b, κ |
| CD49f | A0070 | Human, Mouse | Rat IgG2a, κ |
| CD5 | A0111 | Mouse | Rat IgG2a, κ |
| CD54 | A0074 | Mouse | Rat IgG2b, κ |
| CD55 | A0558 | Mouse | Armenian Hamster IgG |
| CD62E | A0379 | Mouse | Mouse IgG1, κ |

| | | | |
|---|---|---|---|
| CD62L | A0112 | Mouse | Rat IgG2a, κ |
| CD62P | A0229 | Mouse | Mouse IgG2a, κ |
| CD63 | A0559 | Mouse | Rat IgG2a, κ |
| CD64 | A0202 | Mouse | Mouse IgG1, κ |
| CD68 | A0560 | Mouse | Rat IgG2a |
| CD69 | A0197 | Mouse | Armenian Hamster IgG |
| CD71 | A0441 | Mouse | Rat IgG2a, κ |
| CD73 | A0077 | Mouse | Rat IgG1, κ |
| CD79b | A0561 | Mouse | Armenian Hamster IgG |
| CD80 | A0849 | Mouse | Armenian Hamster IgG |
| CD83 | A0562 | Mouse | Rat IgG1, κ |
| CD86 | A0200 | Mouse | Rat IgG2a, κ |
| CD8a | A0002 | Mouse | Rat IgG2a, κ |
| CD8b | A0230 | Mouse | Rat IgG2b, κ |
| CD9 | A0813 | Mouse | Rat IgG2a, κ |
| CD90-1 | A0380 | Rat | Mouse IgG1, κ |
| CD90-2 | A0075 | Mouse | Rat IgG2b, κ |
| CD93 | A0113 | Mouse | Rat IgG2b, κ |
| CD95 | A0917 | Mouse | Mouse IgG1, κ |
| CX3CR1 | A0563 | Mouse | Mouse IgG2a, κ |
| CXCR4 | A0444 | Mouse | Rat IgG2b, κ |
| DLL1 | A0884 | Mouse | Armenian Hamster IgG |
| DoprD4 | A0434 | Human, Mouse | |
| DR3 | A0836 | Mouse | Armenian Hamster IgG |
| ENPP1 | A0891 | Mouse | Rat IgG2b, κ |
| ERK1 | A0222 | Human, Mouse | Rat IgG2a, κ |
| ESAM | A0596 | Mouse | Rat IgG2a, κ |
| F4-80 | A0114 | Mouse | Rat IgG2a, κ |
| FceRIa | A0115 | Mouse | Armenian Hamster IgG |
| FRB | A0564 | Mouse | Rat IgG2a, κ |
| GABRB3 | A0435 | Human, Mouse | |
| IA-IE | A0117 | Mouse | Rat IgG2b, κ |
| IgD | A0571 | Mouse | Rat IgG2a, κ |
| IgG_Hamster | A0241 | Isotype control | Armenian Hamster IgG |
| IgG1_Mouse_ | A0090 | Isotype control | Mouse (BALB/c) IgG1, κ |
| IgG1_Rat_k | A0236 | Isotype control | Rat IgG1, κ |
| IgG1_Rat_l | A0237 | Isotype control | Rat IgG1, λ |
| IgG2a | A0239 | Rat | |
| IgG2a_Mouse | A0091 | Isotype control | Mouse IgG2a, κ |
| IgG2a_Rat_k | A0238 | Isotype control | Rat IgG2a, κ |
| IgG2b_Mouse | A0092 | Isotype control | Mouse IgG2b, κ |
| IgG2b_Rat_k | A0095 | Isotype control | Rat IgG2b, κ |
| IgG2c_Rat_k | A0240 | Isotype control | Rat IgG2c, κ |
| IgM | A0450 | Mouse | Rat IgG2a, κ |
| IL33Ra | A0837 | Mouse | Rat IgG2a, κ |
| Integrin-b7 | A0214 | Human, Mouse | Rat IgG2a, κ |
| IRF4 | A0249 | Human, Mouse | Rat IgG1, κ |
| JAML | A0877 | Mouse | Armenian Hamster IgG |

| | | | |
|---|---|---|---|
| KCC2 | A0438 | Mouse, Rat | |
| KLRG1 | A0250 | Human, Mouse | Syrian Hamster IgG |
| Ly49D | A0841 | Mouse | Rat IgG2a, κ |
| Ly49H | A0839 | Mouse | Mouse IgG1, κ |
| Ly6A-Ly6E | A0130 | Mouse | Rat IgG2a, κ |
| Ly6C | A0013 | Mouse | Rat IgG2c, κ |
| Ly6D | A5106 | Mouse | Rat IgG2c, κ |
| Ly6G | A0015 | Mouse | Rat IgG2a, κ |
| Mac2 | A0895 | Human, Mouse | Rat IgG2a, κ |
| MAdCAM1 | A0232 | Mouse | Rat IgG2a, κ |
| MERTK | A0565 | Mouse | Rat IgG2a, κ |
| NK1-1 | A0118 | Mouse | Mouse IgG2a, κ |
| Notch1 | A0442 | Mouse | Armenian Hamster IgG |
| P2RY12 | A0415 | Mouse | Rat IgG2b, κ |
| P2X7R | A0824 | Mouse | Rat IgG2b, κ |
| Panendotheli | A0381 | Mouse | Rat IgG2a, κ |
| PIRA_PIRB | A0882 | Mouse | Rat IgG1, κ |
| RORg | A0223 | Human, Mouse | Mouse IgG2a, κ |
| SiglecH | A0119 | Mouse | Rat IgG1, κ |
| TCRb | A0120 | Mouse | Armenian Hamster IgG |
| TCRb-V5 | A0354 | Mouse | Mouse IgG1, κ |
| TCRb-V8 | A0235 | Mouse | Rat IgG2a, κ |
| TCRg-V1.1 | A0209 | Mouse | Armenian Hamster IgG |
| TCRg-V2 | A0211 | Mouse | Armenian Hamster IgG |
| TCRg-V3 | A0210 | Mouse | Syrian Hamster IgG |
| TCRgd | A0121 | Mouse | Armenian Hamster IgG |
| TER119 | A0122 | Mouse | Rat IgG2b, κ |
| TIGIT | A0848 | Mouse | Mouse IgG1, κ |
| Tim4 | A0567 | Mouse | Rat IgG2a, κ |
| TLR4 | A0875 | Mouse | Rat IgG2a, κ |
| XCR1 | A0568 | Mouse, Rat | Mouse IgG2b, κ |

| Clone | Gene | Barcode sequence |
|---|---|---|
| TKS-1 | Tnfsf9 | CAGTTCAGTACGCAG |
| 3C4 (MIC2/4) | Icam2 | GATATTCAGTGCGAC |
| 2E7 | Itgae | TTCATTAGCCCGCTG |
| MJ7/18 | Eng | TATCCCTGCCTTGCA |
| 429 (MVCAM.A) | Vcam1 | CGTTCCTACCTACCT |
| 1D4B | Lamp1 | AAATCTGTGCCGTAC |
| AFS98 | Csf1r | TTCCGTTGTTGTGAG |
| 2B8 | Kit | TGCATGTCATCGGTG |
| M17/4 | Itgal | AGAGTCTCCCTTTAG |
| M1/70 | Itgam | TGAAGGCTCATTTGT |
| N418 | Itgax | GTTATGGACGCTTGC |
| 5H4 | Il2rb | GGTATGCGACACTTA |
| I015F8 | Il4ra | GAACCGTAGTATAAC |
| A7R34 | Il7r | GTGTGAGGCACTCTT |
| OX-86 | Tnfrsf4 | CTCACCTACCTATGG |
| A2F10 | Flt3 | GTAGCAAGATTCAAG |
| 17B5 | Tnfrsf9 | TCCCTGTATAGATGA |
| 281-2 | Sdc1 | GCGTTTGTATGTACT |
| Sa14-2 | Cd14 | AACCAACAGTCACGT |
| APA5 | Pdgfra | GTCATTGCGGTCCTA |
| P1H12 | Mcam | CCTTGGATAACATCA |
| MC-480 | Fut4 | GCTAGTTTGTGCTGC |
| TC15-12F12.2 | Slamf1 | CAACGCCTAGAAACC |
| UC10-4B9 | Ctla4 | AGTGTTTGTCCTGGT |
| 93 | Fcgr3/Fcgr2 | TTCGATGCTGGAGCA |
| S15049I | Cd163 | GAGCAAGATTAAGAC |
| 3D6.112 | Siglec1 | ATTGACGACAGTCAT |
| P84 | Sirpa | GATTCCCTTGTAGCA |
| CXCR3-173 | Cxcr3 | GTTCACGCCGTGTTA |
| L138D7 | Cxcr5 | ACGTAGTCACCTAGT |
| 6D5 | Cd19 | ATCAGCCATGTCAGT |
| SA203G11 | Ccr2 | AGTGCGATCTGCAAC |
| J073E5 | Ccr3 | TAGAACCGTATCCGT |
| HM-CCR5 | Ccr5 | ACCAGTTGTCATTAC |
| 29-2L17 | Ccr6 | CTCTCTGCATTCCTC |
| 4B12 | Ccr7 | TTATTAACAGCCCAC |
| 1B1 | Cd1d1 | CAACTTGGCCGAATC |
| RM2-5 | Cd2 | TTGCCGTGTGTTTAA |
| SA275A11 | Ms4a1 | TCCACTCCCTGTATA |
| OX-90 | Cd200 | TCAATTCCGGTAGTC |
| OX-110 | Cd200r1 | ATTCTTTCCCTCTGT |
| Ba13 | Cd200r3 | ATCAACTTGGAGCAG |
| RCR-16 | Procr | TATGATCTGCCCTTG |
| 1F8C33 | Msr1 | AGCTAGACACGTTGT |
| C068C2 | Mrc1 | TCAACTCGGTGTTGC |
| 4C7 | Cd207 | CGATTTGTATTCCCT |
| 7E9 | Cr2/Cr1 | GGATAATTTCGATCC |

| | | |
|---|---|---|
| OX-97 | Cd22 | AGGTCCTCTCTGGAT |
| C9B7W | Lag3 | ATTCCGTCCCTAAGG |
| 10E5 | Cd226 | ACGCAGTATTTCCGA |
| B3B4 | Fcer1g | TCTCTTGGAAGATGA |
| M1/69 | Cd24a | TATATCTTTGCCGCA |
| PC61 | Il2ra | ACCATGAGACACAGT |
| H194-112 | Dpp4 | ATGGCCTGTCATAAT |
| LG.3A10 | Cd27 | CAAGGTATGTCACTG |
| HMHV-1B18 | Tnfrsf14 | GATCCGTGTTGCCTA |
| 6A6 | Btla | TGACCCTATTGAGAA |
| MIH6 | Cd274 | TCGATTCCACCAACT |
| C398.4A | Icos | CGCGCACCCATTAAA |
| RMP1-30 | Pdcd1 | GAAAGTCAAAGCACT |
| 37.51 | Cd28 | ATTAAGAGCGTGTTG |
| HMβ1-1 | Itgb1 | ACGCATTCCTTGTGT |
| 17A2 | Cd3e | GTATGTCCGCTCGAT |
| TX52 | Cd300c | GTGATCTAAGATGCG |
| ZAQ5 | Cd300lg | CGGTCCGTATCATTT |
| LOM-8.7 | Clec10a | TGTATTTACTCACCG |
| URA-1 | Mgl2 | CTTGCCTTGCGATTT |
| 3E12 | Nrp1 | CCAGCTCATTCAACG |
| 89B3A5 | Kdr | AGTTGTCCTGTACGA |
| 390 | Pecam1 | GCTGTAGTATCATGT |
| CX5 | Klrk1 | GAGGCTTATCATTTC |
| 927 | Bst2 | TGTGGTAGCCCTTGT |
| G8.8 | Epcam | ACCCGCGTTAGTATG |
| 29A1.4 | Ncr1 | CCCTTTCACCTCGAA |
| HM34 | Cd34 | GATTCCTTTACGAGC |
| DTA-1 | Tnfrsf18 | GGCACTCTGTAACAT |
| HM36 | Cd36 | TTTGCCGCTACGACA |
| RMT3-23 | Havcr2 | ATTGGCACTCAGATG |
| 7H11 | Clec9a | AACTCAGTTGTGCCG |
| 5D3/CLEC12A | Clec12a | GCGAGAAATCTGCAT |
| 90 | Cd38 | CGTATCCGTCTCCTA |
| Duha59 | Entpd1 | GCGTATTTAACCCGT |
| RM4-5 | Cd4 | AACAAGACCCTTGAG |
| MWReg30 | Itga2b | ACTTGGATGGACACT |
| S11 | Spn | TTGGAGGGTTGTGCT |
| RA3-6B2 | Ptprc | CCTACACCTCATAAT |
| HM48-1 | Cd48 | AGAACCGCCGTAGTT |
| HMα1 | Itga1 | CCATTCATTTGTGGC |
| | Itga2 | CGCGTTAGTAGAGTC |
| R1-2 | Itga4 | CGCTTGGACGCTTAA |
| GoH3 | Itga6 | TTCCGAGGATGATCT |
| 53-7.3 | Cd5 | CAGCTCAGTGTGTTG |
| YN1/1.7.4 | Icam1 | ATAACCGACACAGTG |
| RIKO-3 | Cd55 | ATTGTTGTCAGACCA |
| RME-1/CD62E | Sele | CTCCCTTTGTAACAT |

| | | |
|---|---|---|
| MEL-14 | Sell | TGGGCCTAAGTCATC |
| RMP-1 | Selp | TGTGTGCCGTAGACT |
| NVG-2 | Cd63 | ATCCGACACGTATTA |
| X54-5/7.1 | Fcgr1 | AGCAATTAACGGGAG |
| FA-11 | Cd68 | CTTTCTTTCACGGGA |
| H1.2F3 | Cd69 | TTGTATTCCGCCATT |
| RI7217 | Tfrc | ACCGACCAGTAGACA |
| TY/11.8 | Nt5e | ACACTTAACGTCTGG |
| HM79-12 | Cd79b | TAACTCAGTGCGAGT |
| 16-10A1 | Cd80 | GACCCGGTGTCATTT |
| Michel-19 | Cd83 | TCTCAGGCTTCCTAG |
| GL-1 | Cd86 | CTGGATTTGTGTATC |
| 53-6.7 | Cd8a | TACCCGTAATAGCGT |
| YTS156.7.7 | Cd8b1 | TTCCCTCTATGGAGC |
| MZ3 | Cd9 | TAGCAGTCACTCCTA |
| OX-7 | Thy1 | AGTATGGGATGCAAT |
| 30-H12 | Thy1 | CCGATCAGCCGTTTA |
| AA4.1 | Cd93 | GGTATTTCCTGTGGT |
| SA367H8 | Fas | CACATCGTTTGTGTA |
| SA011F11 | Cx3cr1 | CACTCTCAGTCCTAT |
| L276F12 | Cxcr4 | GTCGTGGTGTTGTTC |
| HMD1-3 | Dll1 | AGACCTCCTTACGAT |
| | Drd4 | TCTCTGCACCGCTTT |
| 4C12 | Tnfrsf25 | GCTTGGGCAATTAAG |
| YE1/19.1 | Enpp1 | CATTAACCGCCCTTA |
| W15133A | Mapk3 | CGCCACTTCATTCAT |
| 1G8/ESAM | Esam | TATAGTTTCCGCCGT |
| BM8 | Adgre1 | TTAACTTCAGCCCGT |
| Mar-01 | Fcer1a | AGTCACCTCGAAGCT |
| 10/FR2 | Folr2 | CTCAGATGCCCTTTA |
| | Gabrb3 | GGTGTGAGCAGTTCT |
| M5/114.15.2 | H2-Eb1 | GGTCACCAGTATGAT |
| 11-26c.2a | Ighd | TCATATCCGTTGTCC |
| HTK888 | / | CCTGTCATTAAGACT |
| MOPC-21 | / | GCCGGACGACATTAA |
| RTK2071 | / | ATCAGATGCCCTCAT |
| G0114F7 | / | GGGAGCGATTCAACT |
| | / | CATTAACAGCGCCAA |
| MOPC-173 | / | CTCCTACCTAAACTG |
| RTK2758 | / | AAGTCAGGTTCGTTT |
| MPC-11 | / | ATATGTATCACGCGA |
| RTK4530 | / | GATTCTTGACGACCT |
| RTK4174 | / | TCCAGGCTAGTCATT |
| RMM-1 | Ighm | AGCTACGCATTCAAT |
| DIH9 | Il1rl1 | GCGATGGAGCATGTT |
| FIB504 | Itgb7 | TCCTTGGATGTACCG |
| IRF4.3E4 | Irf4 | GGATTTGTATCTCCC |
| 4E10 | Jaml | GTTATGGTTCGTGTT |

| | | |
|---|---|---|
| | Slc12a5 | GAGCTTGTACCGCTT |
| 2F1/KLRG1 | Klrg1 | GTAGTAGGCTAGACC |
| 4E5 | Klra4 | TATATCCCTCAACGC |
| 3D10 | Klra8 | CCAGTAGGCTTATTA |
| D7 | Ly6a | TTCCTTTCCTACGCA |
| HK1.4 | Ly6c2 | AAGTCGTGAGGCATG |
| 49-4H | Ly6d | ATGTCCTACCTCAAA |
| 1A8 | Ly6g | ACATTGACGCAACTA |
| M3/38 | Lgals3 | GATGCAATTAGCCGG |
| MECA-367 | Madcam1 | TTGGGCGATTAAGAA |
| 2B10C42 | Mertk | AGTAGAGCAACTCGT |
| PK136 | Klrb1c | GTAACATTACTCGTC |
| HMN1-12 | Notch1 | TCCGGTCACTCAGTA |
| S16007D | P2ry12 | TTGCTTATTTCCGCA |
| 1F11 | P2rx7 | TGCTTCATTCATGTG |
| MECA-32 | Plvap | CGTCCTAGTCATTGG |
| 6C1 | Pirb/Pira1 | TGTAGAGTCAGACCT |
| 2F7-2 | Rorc | TTCCCTACGCCGAAT |
| 551 | Siglech | CCGCACCTACATTAG |
| H57-597 | Tcrb | TCCTATGGGACTCAG |
| MR9-4 | Tcrb | CTCAACAGTATTCTG |
| KJ16-133.18 | Tcrb | ACTATCCGTTGTGCT |
| 2.11 | Tcrb | TCGTTTAACCAGCCT |
| UC3-10A6 | Tcrb | AAGCTGCACCGTAAT |
| 536 | Tcrb | TCGTGGTCCCTTTCT |
| GL3 | Tcrd/Tcrg | AACCCAAATAGCTGA |
| TER-119 | Ly76 | GCGCGTTTGTGCTAT |
| 1G9 | Tigit | GAAAGTCGCCAACAG |
| RMT4-54 | Timd4 | TGCTGGAGGGTATTC |
| MTS510 | Tlr4 | GCAGTTGTCCGATTC |
| ZET | Xcr1 | TCCATTACCCACGTT |
